# Little hope for the polyploid endemic Pyrenean Larkspur (*Delphinium montanum*): evidences from population genomics and Ecological Niche Modelling

**DOI:** 10.1101/2021.09.15.460086

**Authors:** Pascaline Salvado, Pere Aymerich Boixader, Josep Parera, Albert Vila Bonfill, Maria Martin, Céline Quélennec, Jean-Marc Lewin, Valérie Delorme-Hinoux, Joris A. M. Bertrand

## Abstract

Species endemic to restricted geographical ranges represent a particular conservation issue, be it for their heritage interest. In a context of global change, this is particularly the case for plants which belong to high-mountain ecosystems and, because of their ecological requirements, are doomed to survive or disappear on their ‘sky islands’. The Pyrenean Larkspur (*Delphinium montanum*, Ranunculaceae) is endemic to the Eastern part of the Pyrenees (France and Spain). It is now only observable at a dozen of localities and some populations show signs of decline, such as a recurrent lack of flowering. Implementing population genomic approach (e.g. RAD-seq like) is particularly useful to understand genomic patterns of diversity and differentiation in order to provide recommendations in term of conservation. However, it remains challenging for species such as *D. montanum* that are autotetraploid with a large genome size (1C-value > 10 pg) as most methods currently available were developed for diploid species. A Bayesian framework able to call genotypes with uncertainty allowed us to assess genetic diversity and population structure in this system. Our results show evidence for inbreeding (mean *G*_IS_ = 0.361) within all the populations and substantial population structure (mean *G*_ST_ = 0.403) at the metapopulation level. In addition to a lack of connectivity between populations, spatial projections of Ecological Niche Modelling analyses under different climatic scenarios predict a dramatic decrease of suitable habitat for *D. montanum* in the future. Based on these results, we discuss the relevance and feasibility of different conservation measures.

## Introduction

Extinction risk associated to climate change is expected not only to increase but to accelerate for every degree rise in global temperatures (Urban, 2015). In this context, the number of threatened species by reduced population sizes, habitat degradation and habitat fragmentation is increasing. Small and fragmented populations are particularly prone to be affected by drift and inbreeding depression and generally display decreased levels of genetic diversity (e.g. Bergl et al. 2008; Dixo et al. 2009). The increase of anthropogenic and environmental pressures, both being intimately bound, represents an unprecedented threat for vulnerable species. In particular, populations from species of high-mountain ecosystems might not be able to move upward rapidly enough to keep within their thermal tolerances and might be doomed on their ‘sky islands’ (see McCormack et al. 2009, e.g. Kidane et al. 2019, De Gabriel Hernando et al. 2021). Thus, species endemic to specific mountain ranges represent a major conservation issue, if only for their heritage interest. It is critical to characterize the distribution of genetic diversity within- and between-their populations to evaluate the inbreeding risk and study population structure to better understand how isolated they really are, in order to undertake effective conservation actions (Calevo et al. 2021).

The threatened Pyrenean larkspur *Delphinium montanum* DC., 1815 perfectly illustrates this situation. This plant (Family Ranunculaceae), endemic to the Eastern part of the Pyrenees (France and Spain), is now only observable at a dozen of localities (two already got extinct over the 20^th^ century) and its global population size was estimated to range between 7 800 to 10 300 individuals in total (Aymerich et al. 2020). It occurs between alpine and subalpine levels, between 1 600 and 2 400 m (Lopéz-Pujol et al. 2007). *D. montanum* is vulnerable as a result of increasing levels of multiple threats. Some of its populations are located nearby touristic sites (*i.e*. mountain resorts or hiking trails) and are negatively impacted by trampling (Simon et al., 2001). Flowers and fruits predation by herbivores, particularly Pyrenean chamois/isards (*Rupicapra pyrenaica pyrenaica*) was also shown to be responsible for the seed bank’s decrease in some populations (Simon et al. 2001, Aymerich 2003). Importantly, differences in population dynamics are observable from one population to another. Some populations are stable, whereas others show decrease in population size (4-Pedraforca) and/or absence of flowering individuals since 2011 (2-Nohèdes). Pioneering work of Simon et al. (2001) and Lopéz-Pujol et al. (2007) also suggested relatively low effective size as well as, evidence of inbreeding within-populations and significant population structure. However, these two studies were based on a limited number of genetic markers (*i.e*. 7 and 14 allozyme *loci*, respectively) and populations studied (*i.e*. 2 and 7, respectively).

Implementing a population genomic approach on *D. montanum* could be particularly fruitful to inform conservation strategies by providing a substantial increase in the resolution of the genetic marker dataset and genome complexity reduction protocols such as Reduction site-associated DNA sequencing (or RAD-seq, Baird 2008) allow to get such information on tens to hundreds of individuals. However, the features of the genome of the species make *Delphinium montanum* particularly challenging from a technical and methodological point of view. *D. montanum* is an autotetraploid species (Simon et al. 2001; Lopéz-Pujol et al. 2007). Our recent flow cytometry estimates of 10.32 pg/1C for this species (*i.e*. about twice the 1C-value reported for several other diploid *Delphinium* species) corroborates previous knowledge and confirms the large genome size of *D. montanum* (Bertrand et al. unpublished data). A large genome size can be considered has a limitation as itself, be it because in requires higher sequencing efforts and costs to carry out conservation studies. The difficulty to infer genotype and allele frequencies on polyploid species with RAD-seq techniques as well as the restricted set of methods available to analyze such data (see Meirmans and van Tienderen 2013; Dufresne et al. 2014; Meirmans et al. 2018) is also likely to explain the relatively low number of population genomics studies dealing with RAD-seq like techniques on non-model polyploid organisms (Clevenger et al. 2015; van de Peer et al. 2017, Ahmad 2021).

Out the handful of studies that have addressed population genomic questions on polyploid organisms without reference genome, some have considered the genotypes as diploid-like data, an assumption that allows to use classical variant calling tools with default settings (e.g. Brandrud et al. 2017, see also Brandrud et al. 2019; 2020 and Záveská et al. 2019). Consensus allelic information may also be coded in the form of ambiguous sites at heterozygous positions (e.g. Wagner et al. 2020). At last, a couple of methods are able to incorporate ploidy-aware calling algorithms and use genotype likelihoods to infer tetraploid genotypes (e.g. Záveská et al. 2019; Brandrud et al. 2020; Karbstein et al. 2020; Ahmad et al. 2021). In studies for which both kind of methods were used, the latter one generally outputted a higher number of variable sites but signals were found to be overall congruent (Záveská et al. 2019; Brandrud et al. 2020).

The recent studies of Záveská et al. (2019), Brandrud et al. (2020) and Ahmad et al. (2021) used the RAD-seq loci from the catalog generated in the popular pipeline Stacks (Catchen et al. 2011; Catchen et al. 2013) to generate a synthetic reference onto which the raw reads are mapped back before to infer tetraploid genotypes with the approach implemented in EBG (Empirical Bayes Genotyping in Polyploids, Blischak et al. 2018, see also Blischak et al. 2016). As a putative alternative, Clark et al. (2019) proposed a package called PolyRAD to call genotypes with uncertainty from polyploid data in an R environment. PolyRAD can deal with multiple levels of ploidy and is able to use for example population structure, linkage disequilibrium and/or self-fertilization as priors to estimate genotype probabilities. To our best knowledge, this promising and convenient method has been rarely used to implement a population genomic approach on natural populations of polyploid non-model organisms.

In this study, we first aimed at reevaluating the conservation status of *Delphinium montanum* following such population genomics approach. We used a protocol called normalized Genotyping By Sequencing (nGBS) that provided us with thousands of genetic markers (SNPs) from which we inferred patterns of diversity and differentiation with an unprecedented resolution in order to quantify inbreeding and level of connectivity between populations. We then used an Ecological Niche Modelling (ENM) approach to better understand the bioclimatic features that constraint the current geographic distribution of *D. montanum*. Based on several Global Climate Models (CGM), we also predicted the putative evolution of the geographic distribution of the species in the future, at different time periods (2011-2040, 2041-2070 and 2071-2100) and under different, more or less pessimistic but still realistic scenarios of evolution of global warming (or RCPs for Representative Concentration Pathways).

## Materials and Methods

### Geographic distribution and population sampling

The Pyrenean Larskpur (*Delphinium montanum*) geographic distribution ranges from Serra del Cadí-Pedraforca (Fig. 1, localities 4, 5, 8, and 9) up to the Puigmal massif (localities 1, 3 and 6) via Tosa d’Alp massif (locality 7). There, the distance between the most distant sites is about 60 km, and only one locality is located outside this area: Nohèdes (locality 2), in the Madres massif. This latter locality is separated from the Puigmal massif populations by the pit of Cerdagne-Conflent, which could have formed a historical barrier to gene flow. In terms of population size, Cadí-Pedraforca is the most important with up to 70% of the global population size (5 450-6 700 individuals). The sites of the Puigmal massif represent 25 to 30% of the global population (2 200-2 700 individuals). Localities of Madres massif (2) and Tosa d’Alp (7) are numerically low (less than 300, from which none are actually flowering and 35 individuals, respectively).

**Figure 1.**
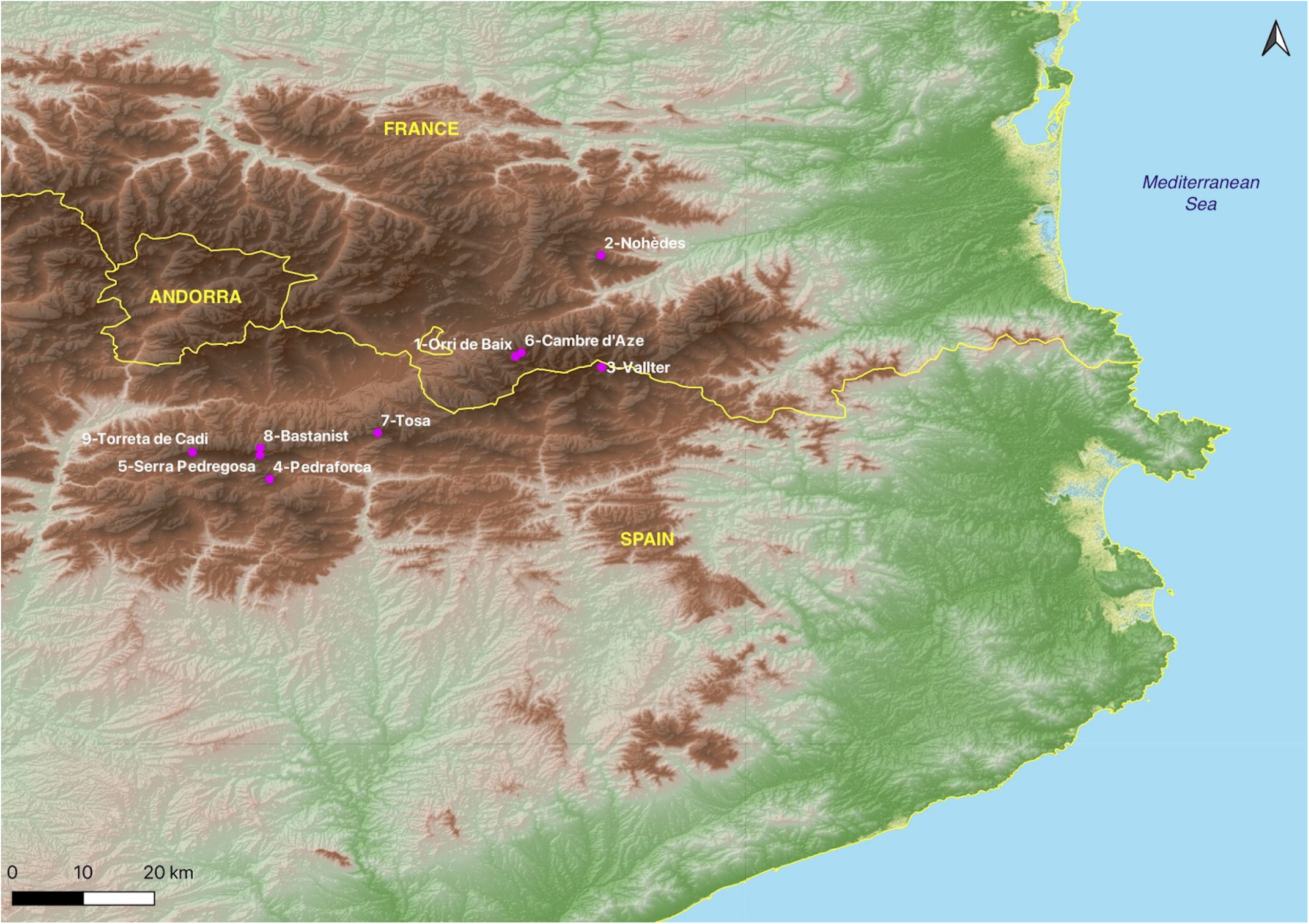
Map representing the sampling localities (pink dots) within the extant geographic distribution of *Delphinium montanum*.

We sampled a total of 106 individuals from 9 localities representative of the geographic distribution of *Delphinium montanum* (in the Eastern Pyrenees: 3 sites in France and 6 in Spain) between July and October 2020 (Table 1, Fig. 1, see also Supplementary Appendix S1). At each locality, we collected on individual distant from at least 1 meter from each other, about 1-2 cm^2^ of leaf tissues that were stored in 90% ethanol at 4°C until DNA extraction. *Delphinium montanum* is not legally protected in France but is considered as vulnerable (VU) on the Red List of threatened species of IUCN France (UICN France, FCBN, AFB, MNHN, 2018). It is legally protected in Catalonia (Decret 172/2008. Catàleg de flora amenaçada de Catalunya) and Spain (Orden AAA/1771/2015. Modificación del catálogo de especies amenazadas). In France, for all sampling done in natural reserves, special authorizations were granted by the ‘Direction Départementale des Territoires et de la Mer 66’ (DDTM 66). In Spain, we asked for and obtained special permits from the Servei de Fauna i Flora de la Generalitat de Catalunya and Natural Parks (Cadí-Moixeró and Ter-Freser).

**Table 1.**
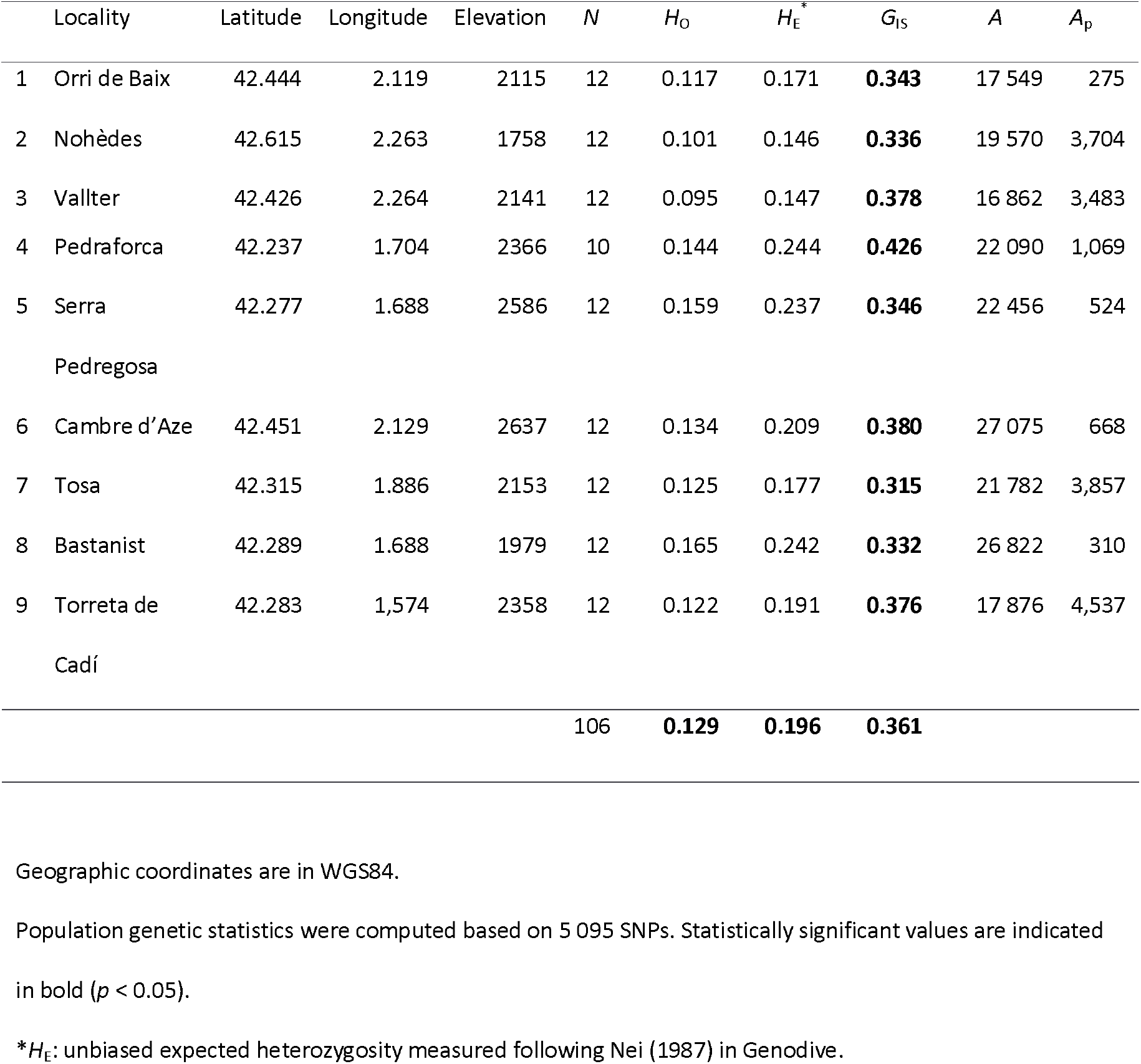
Geographic coordinates (in °) and elevation (in m above sea level) of the sampling localities, sample size (*N*), observed (*H*_O_) and expected (*H*_E_) levels of heterozygosity, deviation from panmixia (*G*_IS_), total number of alleles (*A*) and number of private alleles (*A*_p_) per locality.

### Molecular procedures

Genomic DNA extraction and genotyping were subcontracted to LGC Genomics GmbH (Berlin, Germany). Genotyping-By-Sequencing (GBS) was performed following a specific protocol called nGBS for ‘normalised GBS’. This double digest Restriction site-Associated DNA seq (or RAD-seq) like protocol relies on two restriction enzymes (PstI and ApeKI, in our case) to reduce *D. montanum* genome complexity and includes a normalization step that aims at avoiding repetitive regions. The resulting 106 individually barcoded libraries were sequenced in paired-end mode (2 x 250 bp) on an Illumina NovaSeq 6000 with an expectation of a minimum number of 1.5 million read pairs per sample.

### Genomic data processing

We used Stacks v.2.41 (Catchen et al. 2011; 2013) to build loci from Illumina reads, *de novo* (*i.e*. without aligning reads to a reference genome). Stacks consists of a wrapper of several scripts that are usually run sequentially including: *process_radtags* to demultiplex and clean reads, *denovo_map.pl* to build loci within individuals, create a catalog and match all samples against it and *populations* to further filter the SNPs obtained at the population level and compute basic population genetic statistics. We first optimised several key parameters of the pipeline: *-m* (the minimum number of identical raw reads required to form a putative allele), *-M* (the number of mismatches allowed between alleles to form a locus) and *-n* (the number of mismatches allowed between loci during construction of the catalog) on a subset of 12 individuals representative of the whole data set (*i.e*. geographic origin and coverage). As recommended by several authors, we varied *M* and *n* (fixing *M = n*) while keeping *m* = 3 (see Paris et al. 2017; Rochette & Catchen 2017). The combination *-m* 3, *-M* 3 and *-n* 3 was found to be the most suitable to maximize the number of SNPs, assembled and polymorphic loci in our case and was used to run Stacks on the whole data set (see Supplementary Appendix S2). As *D. montanum* is a tetraploid species, we also specified the option ‘*-X ustacks:--max_locus_stacks* 5’ (in the *ustacks* script’s) to allow the number of alleles per site to reach the number of 5 instead or 3 by default. However, as the *populations* script is unable to deal with polyploid genotypes, we followed a different approach to call variants and complete genomic data processing.

We then used polyRAD (Clark et al. 2019), a method implemented in an R-package to call genotypes with uncertainty from sequencing data in polyploids. The Bayesian genotype caller implemented in polyRAD is able to import read depth from the catalog and matches files from Stacks through a specific function called polyRAD::read_stacks() to further estimate the probability of each possible genotype for each individual and each locus, taking into account features such as possible levels of ploidy and population structure as priors. In order to get similar filtering parameters than with Stacks and following the polyRAD author’s guidelines, we used the following arguments *-min.ind.with.minor. alleles= 5, -min.ind.with.reads= 84* and a ploidy of 4. We then plotted the distribution of the ratio of the individual over expected heterozygosity *H*_ind_/*H*_E_. The peak of the distribution of *H*_ind_/*H*_E_ by locus (0.2) was considered to represent well-behaved markers from which inbreeding (F) was estimated at 0.73. From this estimate of *F*, we then simulated the expected distribution of *H*_ind_/*H*_E_, if all markers were behaving in a Mendelian fashion. Based on the distribution, we defined minimum and maximum thresholds of 0 and 0.406 comprising 95% of the observations and filtered out markers for which *H*_ind_/*H*_E_ were outside this interval. We then tested overdispersion (*i.e*. in our case, how much does read depth distribution deviate from what would be expected under binomial distribution) and adjusted its parameter to 6 (from a range of 2 to 14). This information was used to call genotypes while taking into account population genetic structure. The most likely genotypes of the remaining 5 095 loci were then exported in several formats such as genind objects or Structure (.str).

### Population genomic analyses

To get an overview of the overall genetic diversity and differentiation among individuals and populations, we first performed a Principal Component Analysis (PCA) based on a matrix of 106 individuals (as rows) and 5 095 SNPs (as columns) coded as a genind object with the R-package ‘adegenet’ (Jombart 2008). We then used Genodive v.3.04 (Meirmans 2020) to compute expected and observed heterozygosity (*H*_E_ and *H*_O_, respectively) as well as the number of alleles (*A*) and the number of private alleles was computed with the R-package ‘poppr’ (Kamvar et al. 2014). The deviation from panmixia was evaluated by computing *G*_IS_ and the statistical significance of the obtained values was estimated based on 10 000 permutations, also in Genodive. Overall and pairwise genetic differentiation was also assessed based on *G*-statistics (*G*_ST_, Nei 1987) as implemented in Genodive in a similar manner. Pattern of Isolation By Distance (IBD) was assessed by examining the relationship between linearized *G*_ST_s values (*i.e*. ln(*G*_ST_/(1- *G*_ST_)) and ln-geographical distances (after Rousset 1997) and its statistical significance was tested with a Mantel test between matrices of pairwise *G*_ST_ and geographical distance.

To further investigate population structure and characterize putative migration/admixture event, we used sNMF (Frichot et al. 2014) as implemented in the R-package LEA (Frichot & François 2015) to estimate individual ancestry coefficients based on sparse non-negative matrix factorization algorithms. sNMF is particularly suitable for our genotype dataset as it can deal with polyploid data and is fast enough to be applied on hundreds of individuals and thousands of markers with reasonable computation time. The number of genetic clusters was varied from *K* = 1 to 10, and analyses were run with 10 replicates at each value of *K*.

### Ecological Niche Modelling

In order to assess the future trends of spatial dynamics of *D. montanum* distribution in a context of global change, we conducted Ecological Niche Modelling (ENM) analyses based on known occurrences of the species. Based on 19 bioclimatic layers we downloaded from CHELSA v.2.1 (Karger et al. 2017), we aimed to i) model current environmental niche and ii) predict the spatial evolution of the niche in the future. The bioclimatic layers available in CHELSA are similar to those available in WORLDCLIM (Fick et al. 2017) in a way that they consist of downscaled Global Climate Models (GCM) output temperature and precipitation estimates at a maximal resolution of 30 arc seconds (approximately 1 km). In the current study, we used the most recent version of CHELSA as the methodology and bias correction it implements have been shown to outperforms WORLDCLIM in mountainous environments (Karger et al. 2017; Bobrowski et al. 2021). Current climate data correspond to time averaged variables over the period 1981-2010.

Future climates correspond to estimates from phase 3b of the Inter-Sectoral Impact Model Intercomparison Project (ISIMIP3b) based on output of phase 6 of the Coupled Model Intercomparison Project (CMIP6). They consist of the output of five GCMs (GFDL-ESM4, UKESM1-0-LL, MPI-ESM1-2-HR, IPSL-CM6A-LR and MRI-ESM2-0) and three climate scenario specifiers (ssp126, ssp370, ssp585). Climate scenarios specifiers reflect different Representative Concentration Pathways (RCPs) associated with different radiative forcing values: 2.6, 7.0 and 8.5 W/m^2^ that can be considered as a plausible range of greenhouse gas concentrations that appear to be optimistic, very likely and pessimistic in term of climate change, respectively. Annually averaged data were downloaded for three future time periods: 2011-2040 (“2035”), 2041-2070 (“2055”) and 2071-2100 (“2085”).

The ENM analyses were conducted thanks to the R-package ENMwizard v.0.3.7 (Heming et al. 2018, see also Gutiérrez et al. 2019; Bagley et al. 2020). We first spatially filtered occurrences to keep only those that were at least 1 km away from each other using the R-package ‘spThin’ (Aiello-Lammens et al. 2015). Due to the relatively small spatial scale of the area under study, this resulted in considering a total of 9 out the 106 occurrences initially available. We then examined raw values for the 19 bioclimatic variables extracted at each occurrence point and noticed that the variable bio19 (“precipitation of coldest quarter”) displayed obviously aberrant (*i.e*. negative) values. This variable bio19 was thus discarded from subsequent analyses. From the 18 remaining variables (bio1 to bio18), we selected the less correlated ones (Person correlation coefficient < 0.75) thanks to the R-package ‘caret’ (Kuhn 2019) and kept five variables: bio5 (“mean daily maximum air temperature of the warmest month”), bio7 (“annual range of air temperature”), bio14 (“precipitation amount of the driest month”) and bio15 (“precipitation seasonality”) for further analyses.

The calibration area for the models was created as a buffer of 1.5° around the minimum convex polygon encompassing all occurrences. We used the maximum entropy method (implemented in MaxEnt ver. 3.4.1, Phillips et al. 2006, 2017) to calibrate model and evaluated model performance thanks to the package ENMeval (Muscarella et al. 2014) as implemented in ENMwizard. We evaluated models using a geographic partition scheme of type “block” and optimized two parameters of MaxEnt: the Regularization Multipliers (RM) and the Feature Classes (FC). RM was varied from 0.5 to 4.5, incremented by 0.5 whereas a suite of 15 FCs (L, for Linear, P, for Product, Q, for Quadratic and H for Hinge) or combination of them were evaluated: L, P, Q, H, LP, LQ, LH, PQ, PH, QH, LPQ, LPH, LQH, PQH, LPQH, resulting in a total of 135 models. Model selection was done by computing the corrected Akaike Information Criterion (“LowAIC”) and the Area Under the receiver operating characteristic Curve (“AUC”) in the function ENMWizard::calib_mdl_b(). The relative importance of the different variables was evaluated with the function ENMWizard::get_cont_permimport(). The best selected model was then spatially projected as current niche. For future predictions, a spatial projection of the consensus of the five CGMs was performed for each time period and each RCPs. Area calculations were carried out after applying a 10-percentile (x10ptp) and a Maximum training sensitivity plus specificity (mtss) threshold rules.

## Results

### Genetic diversity and differentiation

We obtained a total of 159 million read pairs across the 106 individuals (2 283 178 to 6 266 452 of raw reads per ind., mean 3 761 722, s.d. = 873 055). We assembled a total of 210 756 loci including 22 911 SNPs with Stacks. Average read depth ranged from 27X to 53X based on the combination of parameters we used. After filtering the data with polyRAD, we finally obtained 32 728 loci including 5 095 SNPs.

The two first PCA axes (PC1 and PC2) represented 14.85% and 9.21% of the total genetic variance, respectively (Fig. 2A). This analysis is consistent with limited amount of within-population differentiation and a clear pattern of between-population differentiation. Overall, populations are arranged according to geography, especially along PC1. In term of genetic diversity, all populations display a significant deviation from panmixia: mean *G*_IS_ =0.361, *p* < 0.05 and population *G*_IS_ values ranging from 0.315 to 0.426 (all *p* < 0.05). However, *G*_IS_ values appear as similar from one population to another (this holds true for the number of total alleles *A*, too, see Table 1). The number of private alleles *A*p is consistent with the relative level of geographic isolation of the populations.

**Figure 2.**
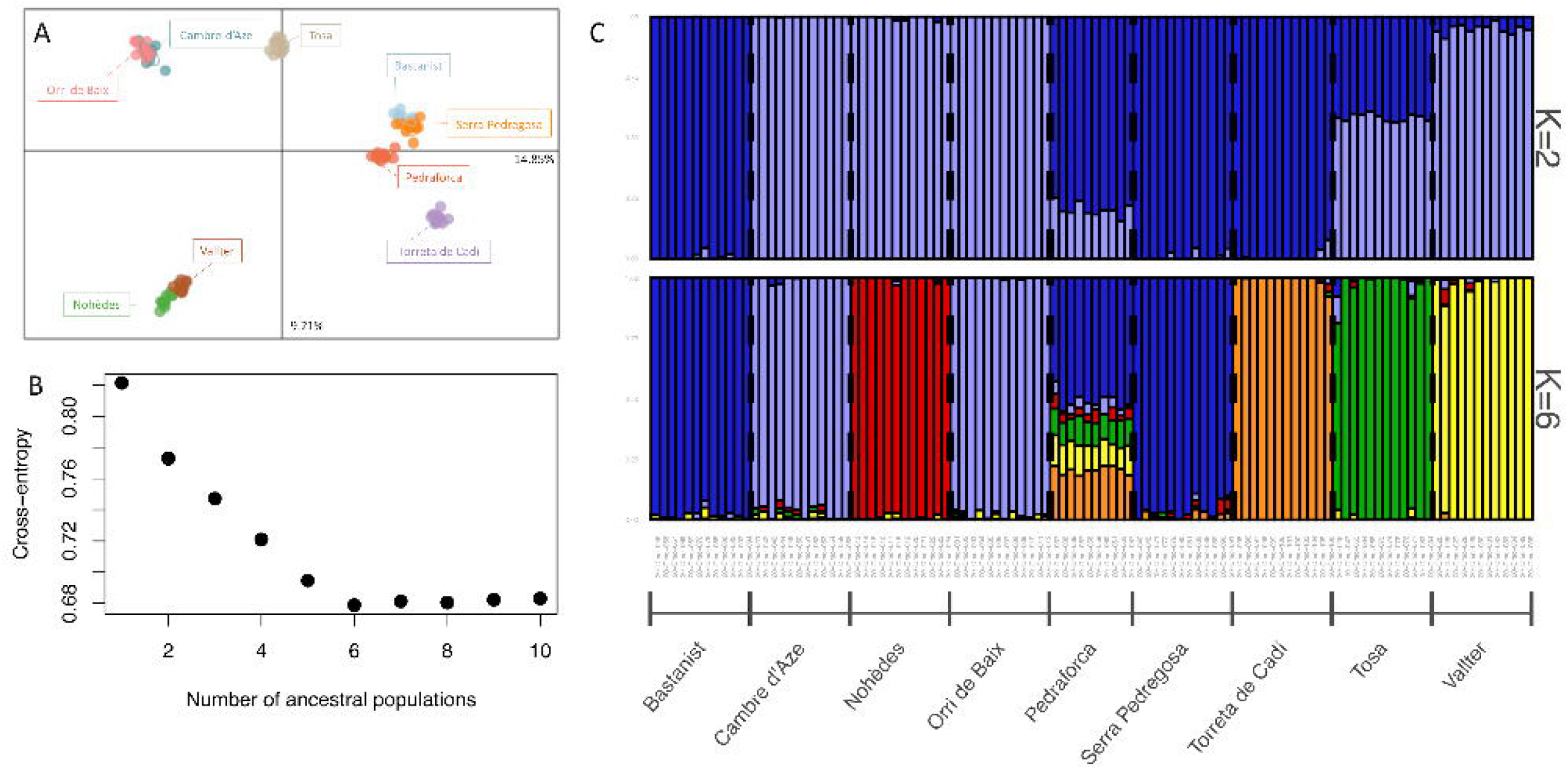
**A)** Principal Component Analysis (PCA) displaying the two first axes (PC1 and PC2) representing 14.85% and 9.21% of the total genetic variance. PCA was computed based on the 106 individuals of *D. montanum* genotyped at 5 095 SNPs whose colours represent sampling localities. B) Values of the cross-entropy criterion for a number of clusters ranging from *K* = 1 to 10 (10 sNMF runs each). The optimal number of *K* was found to be 6. C) Barplot of ancestry coefficients obtained from sNMF for 106 individuals for *K* = 2 and *K* = 6, based on 5 095 SNPs.

The degree of genetic differentiation was found to be relatively high overall (mean *G*_ST_ value = 0.403, *p* < 0.05) and pairwise (with values ranging from 0.156 to 0.527, Supplementary Appendix S3). There is a clear and significant pattern of Isolation By Distance across the dataset (*r* = 0.601, *p* = 0.001, Fig. 3) indicating that gene flow is limited in space at the scale of the *D. montanum* geographic range. The sNMF analysis is also consistent with a strong population structure arranged with geography. Cross-entropy was the lowest for *K*= 6 (Fig. 2B and 2C). Each cluster corresponds to a sampling site with two exceptions. The individuals from the locality of 1-Orri de Baix were grouped together with the close locality of 6-Cambre d’Aze, and the Western localities of the Serra del Cadí (8-Bastanist), 5-Serra Pedregosa and to a lesser extent 4-Pedraforca) formed another group. Only the individuals of 4-Pedraforca showed mixed ancestry coefficients.

**Figure 3.**
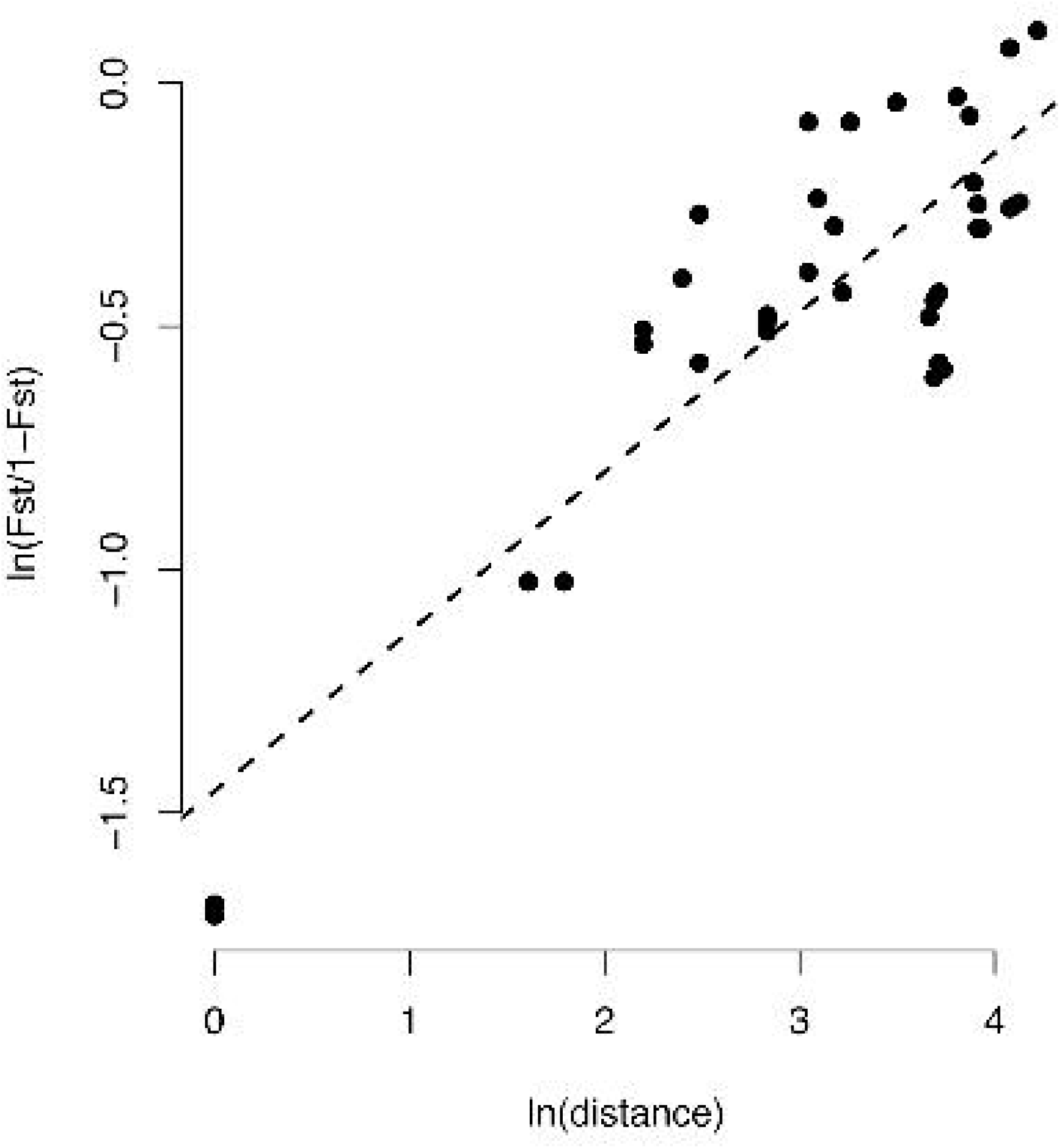
Pattern of Isolation By Distance (IBD) across the data set. Linearized *G*_ST_s values (*i.e*. ln(*G*_ST_/(1- *G*_ST_)) are plotted against ln-geographical distances.

### Suitable habitats for *D. montanum* and prediction of its spatial dynamics

ENMeval analyses identified RM = 2 and a combination of Quadratic and Hinge feature (QH) classes as the best-performing parameters for calibrating the final ENM following the “LowAIC” and M=2 and a combination of Product and Hinge feature (PH) classes following the “AUC” optimality criteria. The first model had the best corrected Akaike Information Criterion (AICc) score (152.25) and an AICc weight of 0.33 whereas the second one had an AICc score of 174.63 and an AICc weight of 4.53.10^-6^. These two models had mean omission rates of 0.375 and 0.25 for the 10^th^ percentile and the lowest presence training thresholds, respectively, as well as a mean test AUC of 0.988. Values of these diagnostic metrics are provided for all candidate models in Supplementary Appendix S4. Both models (RM= 2/PH and RM=2/QH) considered bio15 (“precipitation seasonality”) as being the most important explanatory variable with a contribution of 88.6% and 90.0%, respectively, followed by bio7 (“Temperature annual range (bio5-bio6))” with 6.2% and 7.1% and bio5 “Maximum temperature of the warmest month” with 5.2 and 3.0%; the contribution of bio14 “Precipitation of the driest month”, being 0% with both models. Current suitable area for *D. montanum* was found to be 2 670 and 2 365 km^2^ following “LowAIC” and “AUC” selection model criteria, respectively (the values were identical whatever the threshold method used). All future consensus spatial projections predict a dramatic decrease of the suitable area (Table 2 and Fig. 4). For RCPs 7.0 and 8.5, the suitable area would have completely disappeared for the time period 2071-2100 (but not for RCP 2.6). We also notice some slightly different outcomes depending on the selection model criteria with RCP 8.5 scenarios giving sometimes higher values than RCP 7.0, and even RCP 2.6 (e.g. time period 2011-2040, “Low AIC” criterion).

**Table 2.** Current and future *Delphinium montanum* suitable area (in km^2^) as modelled and predicted from Ecological Niche Modelling under different model selection criteria (LowAIC and AUC) and thresholding methods (10^th^ percentile of training presence, x10ptp and Maximum training sensitivity plus specificity, mtss. Future predictions are reported for several time periods (2011-2040, 2041-2070 and 2071-2100) and various Representative Concentration Pathways (RCP 2.6, 7.0 and 8.5).

| Model selection criterion | Thresholds methods | Current | 2011-2040 |  |  | 2041-2070 |  |  | 2071-2100 |  |  |
| --- | --- | --- | --- | --- | --- | --- | --- | --- | --- | --- | --- |
|  |  |  | RCP 2.6 | RCP 7.0 | RCP 8.5 | RCP 2.6 | RCP 7.0 | RCP 8.5 | RCP 2.6 | RCP 7.0 | RCP 8.5 |
| LowAIC | x10ptp | 2670 | 523 | 492 | 526 | 137 | 17 | 24 | 272 | 0 | 0 |
|  | mtss |  |  |  |  |  |  |  |  |  |  |
| AUC | x10ptp | 2365 | 459 | 455 | 446 | 161 | 25 | 43 | 287 | 0 | 0 |
|  | mtss |  |  |  |  |  |  |  |  |  |  |

**Figure 4.**
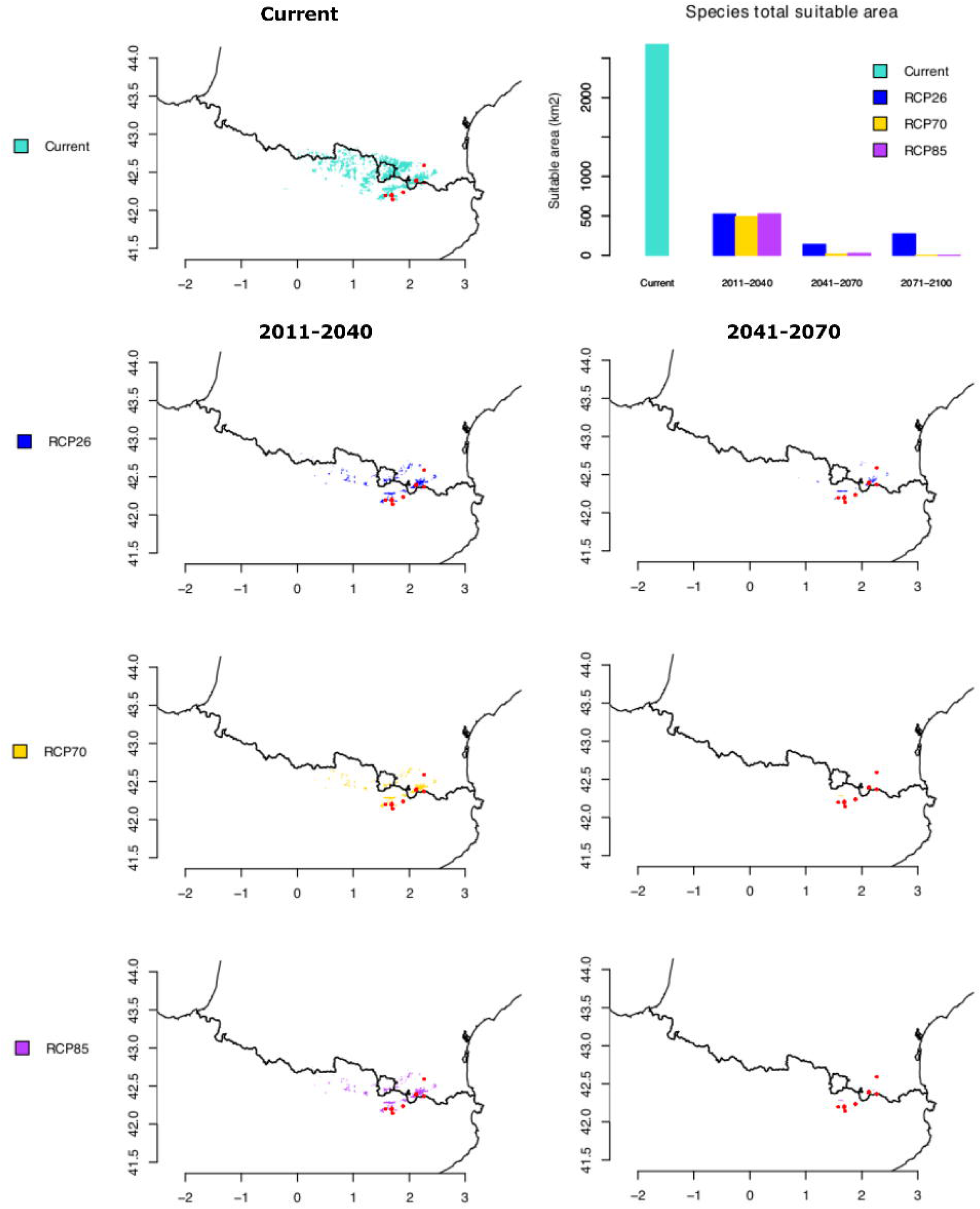
Geographic areas holding suitable climatic conditions for *Delphinium montanum* with projection under present climatic conditions (average for 1981-2010) and consensus projection (from 5 Global Climate Models (CGMs): GFDL-ESM4, UKESM1-0-LL, MPI-ESM1-2-HR, IPSL-CM6A-LR and MRI-ESM2-0), for various plausible climatic scenarios for years 2035 (average for 2011-2040), 2055 (average for 2041-2070) and three Representative Concentration Pathways (RCPs): RCP 26, RCP 70 and RCP 8.5. The time period 2071-2100 is not represented since the suitable area would have completely disappeared for RCPs 7.0 and 8.5. The Barplot displays current and future area values under various plausible climatic scenarios. These values correspond to “LowAIC” model (“AUC” model being very similar).

## Discussion

### Conservation genomics on a polyploid species with large genome size

The use of RAD-seq like (nGBS) data with the method implemented in PolyRAD allowed us to successfully address the challenge of conducting a population genomic approach on a plant species that is both polyploid (autotetraploid) and has a large genome size (> 10 Gbp). It confirms the convenience of the workflow we used, specifically calling genotypes in an R-environment, as an alternative to the ones followed in the handful of studies recently published on similar systems (e.g. Záveská et al. 2019; Brandrud et al. 2020; Karbstein et al. 2020; Ahmad et al. 2021). From the > 5 000 SNPs we could rely on, we were able to get an unprecedented genomic picture of the diversity and the differentiation of the species in order to propose appropriate conservation recommendations. The results we obtained at the genome scale are overall consistent with previous knowledge on population genetics of *D. montanum* (see Simon et al. 2001; López-Pujol 2007).

### A set of populations isolated on sky islands

Our results confirm that populations of *D. montanum* can be seen as a set of island-like dwelling systems separated from each other by a matrix of unsuitable habitats and that gene flow has little to no chance to be efficiently maintained within this metapopulation. All populations considered in this study were found to display deficit in heterozygotes (*G*_IS_ > 0.315, *p* < 0.05). These results confirm previous findings of Simon et al. (2001) and López-Pujol et al. (2007) and show that the autotetraploid status of the species does not impede relatively low levels of heterozygosity (*H*_O_ < 0.17). Polyploidy is expected to be associated with increased genetic variation, a characteristic that provides more diversity to adapt (especially to extreme environments such as mountainous ones) and allows buffering effect against deleterious mutations (see van de Peer et al. 2021 and references therein). From a conservation point of view, polyploidy does not seem to be associated with increased risk of extinction even though plant with big genome size may be more prone to be under threat (Vinogradov 2003; Pandit et al. 2011). As two other closely related congeneric species of *Delphinium montanum*: *D. dubium* and *D. oxysepalum*, that are endemic to the Alps and the Carpathians, respectively, are also tetraploid (López-Pujol et al. 2007), we speculate that polyploidy may have allowed to these species to adapt and survive into these mountainous environments. The open question is whether current levels of genetic diversity may be sufficient or not to allow *D. montanum* populations to overcome inbreeding depression and provide sufficient variation for adaptation in a context of global change. It should be noted that levels of genetic diversity are overall similar among the populations investigated and we could not associate obvious signs of decline of some of them (e.g. Nohèdes, Pedraforca) to decreased levels in genetic diversity or evidence for deviation from panmixia. This suggests that the evidences of possible decrease in effective size may be too recent to be detectable at the genetic level (see Peery et al. 2012). Unfortunately, the limited number of available methods for polyploids makes effective size estimation and test for bottleneck events cumbersome to perform in such systems.

The PCA, the clustering analyses, the *F*_ST_ values as well as the pattern of Isolation By Distance, further confirm limited gene flow between populations and the existence of a population structure that is consistent with geography. The most likely number of clusters found in the sNMF analysis (*K*=6) globally distinguishes populations from the Serra del Cadí (8-Bastanist, 5-Serra Pedregosa and 4-Pedraforca) from those of the Puigmal range (1-Orri de Baix and 6-Cambre d’Aze) plus the satellite populations of 3-Vallter, 2-Nohèdes, 7-Tosa d’Alp and 9-Torreta de Cadí (which is the westernmost locality, a bit distant from the others in the Serra del Cadí). Only the locality of 4-Pedraforca shows some ambiguity in population assignment that could indicate a mixed origin. At *K*=2, the intermediate status of 7-Tosa d’Alp (and to a lesser extent of 4-Pedraforca) are also in agreement with its intermediate geographic position and may reflect an ancestral split between Northern and Southern populations. Altogether, these results support that the insect pollinators (mainly bumblebees) and the natural dispersal of seeds produced (barochoric) are not able to maintain gene flow over the geographic distances involved.

### Climate change as the main threat for *Delphinium montanum*

The two previous studies of Simon et al. (2001) and López-Pujol et al. (2007) have listed several threats to explain the decline of *D. montanum*, at least in some particular populations. According to Simon et al. (2001), natural disturbance of habitats due to rock falls and avalanches, trampling because of an overall increase in human visits and animal predation (e.g. caterpillar, Pyrenean chamois/isards and perhaps red and roe deers) as well as competition for insect pollinators with simultaneously blooming species (e.g. *Aconitum* spp.) have thus been considered as more or less serious problems for *D. montanum*. The most recent study of López-Pujol et al. (2007) also emphasized on the consequences of the erosion of genetic diversity, especially the loss of rare alleles associated with weak possibility of gene flow between localities that could be however compensated by relatively high effective size in several populations. Without ruling out all these threats on *D. montanum* populations, our results rather support climate change as the main threat for this species. The future predictions of the area of habitat suitability all agree that the spatial extent of the ecological niche of *D. montanum* is very likely to dramatically decrease, and even disappear by the end of the 21^th^ century, with some differences based on the various scenarios we considered. The current suitable habitat for *D. montanum* has an area that was estimated at about 2500 km^2^, an area that would decrease to about 500 km^2^ (*i.e*. −75 %), by 2040 whatever the scenario envisaged. By 2070, this area would again have decreased to about 150 km^2^ based on RCP 2.6 and up to less than 50 km^2^ based on the two other scenarios (RCP 7 and RCP 8.5). A Representative Concentration Pathway of 2.6 W/m^2^ has to be considered as a rather optimistic pathway (that would keep global temperature rise below 2°C by 2100). It would require that CO_2_ emissions start declining by 2020 and go to zero by 2100 (as well as declines in emission of other greenhouses gases, such as CH_4_, SO_2_). RCP 7 is usually considered as a baseline rather than a mitigation target. According to RCP 8.5, emissions would continue to rise throughout the 21^st^ century and can be considered as a pessimistic scenario (with an increase in average temperature in the order of 4°C by 2100). The various output of these scenarios likely explains why the bioclimatic niche of *D. montanum* would have completely disappeared by 2100 for RCP 7 and 8.5 but could have persisted and even started to increase again to about 10% of its current suitable area following the RCP 2.6 scenario.

Environmental changes as the main threat for *D. montanum* populations is also supported from field observations. The population of Nohèdes (2) have not flowered since 2011. The individuals there are very small (*i.e*. a few centimetres high) and remain at a vegetative stage. On this site, we could not notice any evidence of avalanche or landslide that could have altered the habitat. In addition, other plants species present in the vicinity (e.g. *Orchis spitzelii, Maianthemum bifolium* and *Convallaria majalis*) seem to suffer from flowering default, further supporting that this phenomenon could be non-specific to *D. montanum* and perhaps caused by bioclimatic or at least environmental modifications. There, output from Regional Climate Models (RCMs) predict an increase in mean annual temperature that could reach 3.1 to 4.5°C (+1.5°C already observed since the mean of the period 1961-1990) associated with a decrease in precipitation of up to 25% and perhaps more compared to present-day conditions (Lespinas et al. 2014). According to the same study, precipitations would mostly decrease in summer (−40%) and in spring (−20%) but would remain similar the rest of the year. This in accordance with our ENM predictions suggesting that bio15 “Precipitation seasonality”, the annual coefficient of variation of rainfall may be by far, the most important variable in explaining the bioclimatic niche of *D. montanum*.

### Recommendations for conservation

*Ex-situ* conservation would be a way to preserve the genetic diversity of *D. montanum*, but also a bridge for future *in-situ* conservation efforts. A seed collection of different populations of *D. montanum* could be stored in a bank. However, this measure would be of limited interest if the decline’s cause of *D. montanum* is actually climate change as keeping dormant seeds will probably not allow this species to adapt to future global changes. Alternatively, conserving genetic diversity in a botanical garden may be a good option to test whether and if so, leave time to plants to adapt to rising temperatures and rainfall modifications. Nevertheless, this approach has potential drawbacks like artificial selection and habitat conversion if *D. montanum* adapts to *ex-situ* environment, which may differ from their initial natural environment (Ren et al. 2014). Moreover, if, growing individuals from the all 9 localities studied in an appropriate botanical garden could be a way to set-up *in-situ* conservation, preliminary experiments have showed that *cultivating D. montanum* under experimental conditions in a greenhouse is very difficult because of its environmental requirements (Aymerich, pers. comm.). This confirms that if any, botanical garden would have to be set up at a relatively high elevation (*i.e*. > 1500 m), where environmental conditions are similar to the ecological optimum of the species.

We can identify three main methods of *in-situ* conservation that could be envisaged for *D. montanum* (see Godefroid et al. 2011; Ren et al. 2014; Mashinski & Albrecht 2017): reinforcement, translocation and managed relocation (or assisted migration). The first one is reinforcement, which consists in adding *ex-situ* grown individuals in their natural population of origin to increase the total number of individuals, increase the spatial occupancy of the species and thereby reduce extinction risk. On the one hand, the increase in population size provided by reinforcement could contribute to counteract the effect of genetic drift. On the other hand, it would have a limited impact to impede inbreeding as all the individuals would originate from the same genetic pool. The second method consists in translocating individuals (seeds, seedlings or adult plants) from a population to another one with the same ecogeography, to increase genetic diversity (the so-called “genetic rescue”) and thus increase the likelihood of population persistence in a changing climate (Mashinski and Albrecht 2017). Our results show that *D. montanum* populations harbor substantial levels of pairwise genetic differentiation which suggest that they have exchanged few or no genetic migrants for many generations and have possibly evolved distinct adaptative differences. As a consequence, gene pool mixing may not be appropriate, since the risk of outbreeding depression could be non-negligible between individuals from the different part of the geographic distribution of *D. montanum*. It could be thus careful to first experiment *ex-situ* crossings to check if hybrids are viable, and do not suffer from outbreeding depression. However, the genetic benefits of mixing source populations during genetic rescue have been shown to generally outweigh the risks of outbreeding depression and loss of local adaptation (see Whiteley et al. 2015; Frankham 2015; Ralls et al. 2018, see also Liddell 2021). For example, translocations with mixed source populations of *Arnica montana* showed that genetic variation remained similar or increased across successive environments and some authors documented a 10-fold increase in population size following gene flow between wild Trinidadian guppy populations (Albretch and Edwards 2020; Fitzpatrick et al. 2020).

Finally, the third method consists in introducing a new population in a suitable new ecosystem, where *D. montanum* has not been observed yet. This method is called managed relocation or assisted migration (Mashinski and Albrecht 2017). However, it may represent a risk of modifying the ecosystem, by creating competition for space and pollinators with other plants naturally present, or by introducing new pathogens. Thus, these three methods of reinforcement, translocation, and managed relocation can present risks of outbreeding depression and ecosystem destabilization, and their benefits may not outweigh those risks. Moreover, since the main threat for *D. montanum* is climate change, on a longer-scale, all those efforts could be unsuccessful. Keeping individuals in a botanical garden seems to be the least harmful and risky method for natural populations today and it could be a good start for crossing experiments between populations.

For the very peculiar situation of 2-Nohèdes’s locality, which seems to be the population the most at risk of extinction at the moment, there is still the question of the lack of flowering. To test whether this problem may be associated with the absence of a specific environmental stimulus, we may propose to either i) experimentally modify temperature/water supply, *in situ*, ii) translocate some individuals to localities where *D. montanum* seem healthy (e.g. Cadí range) and see if flowering can be restored and/or iii) experimentally modify temperature/water supply in the lab to better understand the conditions required for flowering and figure out the limiting factor in the wild. Alternatively, it may be due to the fact that that this locality is formed by immature individuals and that the local population itself may be recovering from an important mortality event. At the moment, we have no further arguments to support this scenario.

### Conclusion

Overall, our results confirm that the remaining populations of *D. montanum* show evidence of decreased genetic diversity and evidence of inbreeding at a genomic scale. The degree of genetic structure observable is consistent with a set of strongly isolated populations no longer able to maintain gene flow between their sky islands. In this context, *in-situ* conservation methods are likely to have an effect on the status of the species. Reinforcement would help counteracting decrease in local population sizes, and translocation would help increasing genetic diversity, buffering inbreeding depression and perhaps, bring the variation required for *D. montanum* to adapt. This being said, *ex-situ* crossings are first recommended to verify the absence of outbreeding depression as local genetic pools were found to differ substantially. As our results support that environmental change may represent the main risk of extinction for *D. montanum* all these measures may be however considered as of limited interest on the long term, if the ecological niche of the species disappears.

## Acknowledgements

This study was funded by the interreg-POCTEFA program (FLORALAB Project EFA294/19) and is set within the framework of the “Laboratoires d’Excellences (LABEX)” TULIP (ANR-10-LBX-41) as well as within the framework “École Universitaire de Recherche (EUR)” TULIP-GS (ANR-18-EURE-0019). We thank Xavier Oliver Martínez-Fornés, Sandra Mendez, Kimberley Goudédranche, Lucie Schaad, Irene Abad, Queralt Tor and all other contributors from the FLORALAB network for the help provided in the field. We also thank Olivier François, Lindsey Clark, Patrick Meirmans and Neander Marcel Heming for technical advice and insightful discussions.

## Authors’ Contributions and Conflict of Interest

PS, VH, JAMB analysed the data and wrote the manuscript. PAB, JP, AVB, MM, CC, JML and JAMB contributed to fieldwork. MM, CC and VH coordinated the FLORALAB network and/or project aspects related to this work and JAMB contributed to the conception of the study. All authors have read and approved the manuscript. The authors declare no conflict of interest.

## Data Accessibility Statement

Sequencing data have been submitted to the European Nucleotide Archive (ENA; https://www.ebi.ac.uk.ena) under Study with accession n°PRJEB46773 and samples accession n°ERS7180281 (SAMEA9457265) and n°ERS7643565 (SAMEA9965271) to n°ERS7643669 (SAMEA9965375).

## References

Ahmad, M., Leroy, T., Krigas, N., Temsch, E.M., Weiss-Scheeweiss, H., Lexer, C., Sehr, E.M., Paun, O. (2021) Spatial and ecological drivers of population structure in *Alkanna tinctiria* (Boraginaceae), a polyploid medicinal herb. Frontiers in Plant Science, 12:706574.

Aiello-Lammens, M.E., Boria, R.A., Radosavljevic, A., Vilela, B., Anderson, R.P. (2015) spThin: an R package for spatial thinning of species occurrence records for use in ecological niche models. Ecography, 38, 541–545.

Albrecht, M.A., Edwards, C.E. (2020) Genetic monitoring to assess the success of restoring rare plant populations with mixed gene pools. Molecular Ecology, 29, 4037–4039.

Aymerich, P. (2003) Effects of wild ungulates predation in the conservation of rare plants: two study cases of the Eastern Pyrenees. Acta Botanica Barcinonensia, 49, 147–164.

Aymerich, P., Martínez-Fornés, X.O., Mendez, S., Mangeot, A., Martin, M., Tenas, B. (2019) Seguiment de l’endemisme dels Pireneus orientals *Delphinium montanum* per la xarva transfronterera FloraCat, Actes del XII Colloqui Internacional de Botànica Pirenaica - Cantàbrica, 147–165.

Bagley, J.C., Heming, N.M., Gutiérrez, E.E., Devisetty, U.K., Mock, K.E., Eckert, A.J., Strauss, S.H. (2020) Genotyping-by-sequencing and ecological niche modeling illuminate phylogeography, admixture, and Pleistocene range dynamics in quaking aspen (*Populus tremuloides*). Ecology and Evolution, 10, 4609–4629.

Baird, N.A., Etter, P.D., Atwood, T.S., Currey, M.C., Shiver, A.L., Lewis, Z.A., Selker, E.U., Cresko, W.A., Johnson, E.A. (2008) Rapid SNP discovery and genetic mapping using sequenced RAD markers. PloS ONE, 3: e3376.

Bergl, R.A., Bradley, B.J., Nsubuga, A., Vigilant, L. (2008) Effects of habitat fragmentation, population size and demographic history on genetic diversity: The Cross River Gorilla in a comparative context. American Journal of Primatology, 70, 848–859.

Blischak, P.D., Kubatko, L.S., Wolfe, A.D. (2016) Accounting for genotype uncertainty in the estimation of allele frequencies in autopolyploids. Molecular Ecology Resources. 16, 742–754.

Blischak, P.D., Kubatko, L.S., Wolfe, A.D. (2018) SNP genotyping and parameter estimation in polyploids using low-coverage sequencing data. Bioinformatics, 34, 407–415.

Bobrowski, M., Weidinger, J., Schickfoff, U. (2021) Is new always better? Frontiers in global climate datasets for modelling treeline species in the Himalayas. Atmosphere 12: 543.

Brandrud, M.K., Paun, O., Lorenzo, M.T., Nordal, I., Brysting, A.K. (2017) RADseq provides evidence for parallel ecotypic divergence in the autotetraploid *Cochlearia officinalis* in Northern Norway. Scientific Reports, 7: 5573.

Brandrud, M.K., Paun, O., Lorenz, R., Baar, J., Hedrén, M. (2019) Restriction-site associated DNA sequencing of *Nigritella* and *Gymnadenia* (Orchidaceae). Molecular Phylogenetics and Evolution, 136, 21–28.

Brandrud, M.K., Baar, J., Lorenzo, M.T., Athanasiadis, A., Bateman, R.M., Chase, M.W., Hedrén, M., Paun, O. (2020). Phylogenomic relationships of diploids and the origins of allotetraploids in *Dactylorhiza* (Orchidaceae). Systematic Biology, 69, 91–109.

Calevo, J., Gargiulo, R., Bersweden, L., Viruel, J., González-Montelongo, C., Rebbas, K., Boutafia L, Fay, M.F. (2021) Molecular evidence of species- and subspecies-level distinctions in the rare *Orchis patens s.l*. and implications for conservation. Biodiversity and Conservation, 30, 1293–1314.

Catchen, J., Amores, A., Hohenlohe, P., Cresko, W., Postlethwait, J. (2011) Stacks: building and genotyping loci de novo from short-read sequences. G3 – Genes, Genomes, Genetics, 1, 171–182.

Catchen, J., Hohenlohe, P.A., Bassham, S., Amores, A., Cresko, W.A. (2013) Stacks: an analysis tool set for population genomics. Molecular Ecology, 22, 3124–3140.

Clark, L.V., Lipka, A.E., Sacks, E.J. (2019) polyRAD: Genotype calling with uncertainty from sequencing data in polyploids and diploids. G3 – Genes, Genomes, Genetics, 9, 663–673.

Clevenger, J., Chavarro, C., Pearl, S.A., Ozias-Akins, P., Jackson, S.A. (2015) Single Nucleotide Polymorphism identification in polyploids: a review, example, and recommendations. Molecular Plant, 8, 831–846.

De Gabriel Hernando, M., Fernandéz-Gil, J., Roa, I., Juan, J., Ortega, F., De la Calzada, F., Revilla, E. (2021) Warming threatens habitat suitability and breeding occupancy of rear-edge bird specialists. Ecography, 44, 1191–1204.

Dixo, M., Metzger, J.P., Morgante, J.S., Zamudio, K.R. (2009) Habitat fragmentation reduces genetic diversity and connectivity among toad populations in the Brazilian Atlantic Coastal Forest. Biological Conservation, 142, 1560–1569.

Dufresne, F., Stift, M., Vergilino, R., Mable, B.K. (2014) Recent progress and challenges in population genetics of polyploid organisms: an overview of current state-of-the-art molecular and statistical tools. Molecular Ecology. 23, 40–69.

Fick, S.E., Hijmans, R.J. (2017) New 1 km spatial resolution climate surfaces for global land areas. International Journal of Climatology, 37, 4302–4315.

Fitzpatrick, S.W., Bradburd, G.S., Kremer, C.T., Salerno, P.E., Angeloni, L.M., Funk, W.C. (2020) Genomic and fitness consequences of genetic rescue in wild populations. Current Biology, 30, 517–522.

Frankham, R. (2015) Genetic rescue of small inbred populations: meta-analysis reveals large and consistent benefits of gene flow. Molecular Ecology, 24, 2610–2618.

Frichot, E., Mathieu, F., Trouillon, T., Bouchard, G., François, O. (2014) Fast and efficient estimation of individual ancestry coefficients. Genetics 4, 973–983.

Frichot, E., François, O. (2015) LEA: An R package for landscape and ecological association studies. Methods in Ecology and Evolution, 6, 925–929.

Godefroid, S., Piazza, C., Graziano, R., Buord, S., Stevens, A.-D., Aguraiuja, R., Cowell, C., Weekley, C.W., Vogg, G., Iriondo, J.M., Johnson, I., Dixon, B., Gordon, D., Magnanon, S., Valentin, B., Bjureke, K., Koopman, R., Vicens, M., Venderborght, T. (2011) How successful are plant species reintroductions? Biological Conservation, 144, 672–682.

Gutiérrez, E.E., Heming, N.M., Penido, G., Dalponte, J.C., Lacerda, A.C.R., Moratelli, R., de Moura Bubadué, J., da Silva, L.H., Wolf, M.M., Marinho-Filho, J. (2019) Climate change and its potential impact on the conservation of the Hoary Fox, *Lycalopex vetulus* (Mammalia: Canidae). Mammalian Biology, 98, 91–101.

Ren, H., Jian, S., Liu, H., Zhang, Q., Lu, F. (2014) Advances in the reintroduction of rare and endangered wild plant species. Science China Life Sciences, 57, 603–609.

Heming, N.M., Dambros, C., Gutiérrez, E. (2018) ENMwizard: advanced techniques for Ecological Niche Modeling made easy. https://github.com/HemingNM/ENMwizard

Jombart, T. (2008) adegenet: a R package for the multivariate analysis of genetic markers. Bioinformatics, 24, 1403–1405.

Kamvar, Z.N., Tabima, J.F., Grünwald, N.J. (2014) Poppr: an R package for genetic analysis of populations with clonal, partially clonal, and/or sexual reproduction. PeerJ, 2: e281.

Karbstein, K., Tomasello, S., Hodač, L., Dunkel, F.G., Daubert, M., Hörandl, E. (2020) Phylogenomics supported by geometric morphometrics reveals delimitation of sexual species within the polyploid apomictic *Ranunculus auricomus* complex (Ranunculaceae). Taxon, 69, 1191–1220.

Karger, D.N., Conrad, O., Böher, J., Kawohl, T., Kreft, H., Soria-Auza, R.W., Zimmermann, N.E., Linder, H.P., Kessler, M. (2017). Climatologies at high resolution for the earth’s land surface areas. Scientific Data, 4: 170122.

Kidane, Y.O., Steinbauer, M.J., Beierkuhnlein, C. (2019) Dead end for endemic plant species? A biodiversity hotspot under pressure. Global Ecology and Conservation, 19: e00670.

Kuhn, M. (2019) Caret: classification and regression training. https://CRAN.R-project.org/package=caret.

Liddell, E., Sunnucks, P., Cook, C.N. (2021) To mix or not to mix gene pools for threatened species management? Few studies use genetic data to examine the risks of both actions, but failing to do so leads disproportionately to recommendations for separate management. Biological Conservation, 256:109072.

Lespinas, F., Ludwig, W., Heussner, S. (2014) Hydrological and climatic uncertainties associated with modeling the impact of climate change on water resources of small Mediterranean coastal rivers. Journal of Hydrology, 511, 403–422.

López-Pujol, J., Orellana, M. R., Bosch, M., Simon, J., Blanché, C. (2007) Low genetic diversity and allozymic evidence for autopolyploidy in the tetraploid Pyrenean endemic larkspur *Delphinium montanum* (Ranunculaceae). Biological Journal of the Linnean Society, 155, 211–222.

Maschinski, J., Albretcht, M.A. (2017) Center for plant conservation’s best practice guidelines for the reintroduction of rare plants. Plant Diversity, 39, 390–395.

McCormack, J.E., Huang, H., Knowles, L.L. (2009) Sky Islands. In: Gillespie, R.G., Clague, D. (eds) Encyclopedia of Islands, University of California Press: Berkeley, CA Pp 839–843.

Meirmans, P.G., Van Tienderen, P.H. (2013) The effects of inheritance in tetraploids on genetic diversity and population divergence. Heredity, 110, 131–137.

Meirmans, P.G., Liu, S., van Tienderen, P.H. (2018) The analysis of polyploid genetic data. Journal of Heredity, 109, 283–296.

Meirmans, P.G. (2020) Genodive version 3.0: Easy-to-use software for the analysis of genetic data of diploids and polyploids. Molecular Ecology Resources, 20, 1126–1131.

Muscarella, R., Galante, P.J., Soley-Guardia, M., Boria, R.A., Kass, J.M., Uriarte, M., Anderson, R.P. (2014) ENMeval: an R package for conducting spatially independent evaluations and estimating optimal model complexity for Maxent ecological niche models. Methods in Ecology and Evolution, 5, 1198–1205.

Nei, M. (1987) Molecular Evolutionary Genetics. Columbia University Press: New York: 1–514.

Pandit, M.K., Pocock, M.J.O., Kunin, W.E. (2011) Ploidy influences rarity and invasiveness in plants. Journal of Ecology, 99, 1108–115.

Paris, J.R., Stevens, J.R., Catchen, J. (2017) Lost in parameter space: a road map for stacks. Methods in Ecology and Evolution, 8, 1360–1373.

Peery, M.Z., Kirby, R., Reid, B.N., Stoelting, R., Doucet-Bëer, E., Robinson, S., Vásquez-Carillo, C., Pauli, J.N., Palsbøll, P. J. (2012) Reliability of genetic bottleneck tests for detecting recent population declines. Molecular Ecology, 21, 3403–3418.

Phillips, S.J., Anderson, R.P., Schapire, R.E. (2006) Maximum entropy modelling of species geographic distributions. Ecological Modelling, 190, 231–259.

Philipps, S.J., Anderson, R.P., Dudík, M., Shapire, R.E., Blair, M.E. (2017) Opening the black box: an open-source release of Maxent. Ecography, 40, 887–893.

Ralls, K., Ballou, J.D., Dudash, M.R., Eldridge, M.D.B., Fenster, C.B., Lacy, R.C., Sunnucks, P. & Frankham, R. (2018) Call for a paradigm shift in the genetic management of fragmented populations. Conservation Letters, 11: e12412.

Ren, H., Jian, S.G., Liu, H.W., Zhang, Q.M., Lu, H.F. (2014) Advances in the reintroduction of rare and endangered wild plant species. Science China Life Sciences, 57, 603–609.

Rochette, N.C., Catchen, J.M. (2017) Deriving genotypes from RAD-seq short read data using Stacks. Nature Protocols, 12, 2640–2659.

Rousset, F. (1997) Genetic differentiation and estimation of gene flow from F-statistics under Isolation by Distance. Genetics, 145, 1219–1228.

Simon, J., Bosch, M., Molero, J., Blanché, C. (2001) Conservation biology of the Pyrenean larkspur (*Delphinium montanum):* a case of conflict of plant versus animal conservation? Biological Conservation, 98, 305–314.

UICN France, FCBN, AFB, MNHN (2018) La liste rouge des espèces menacées en France – Chapitre Flore vasculaire de France métropolitaine. Paris, France.

Urban, M.C. (2015) Accelerating extinction risk from climate change. Science, 348, 571–573.

Van de Peer, Y., Mizrachi, E., Marchal, K. (2017) The evolutionary significance of polyploidy. Nature Review Genetics, 18, 411–424.

Van de Peer, Y., Ahsman, T.-L., Soltis, P.S., Soltis, D.E. (2021) Polyploidy: an evolutionary and ecological force in stressful times. Plant Cell, 33, 11–26.

Vinogradov, A.E. (2003) Selfish DNA is maladaptive: evidence from the plant Red List. Trends inGenetics, 19, 609–614.

Wagner, N.D., He, L., Hörandl, E. (2021) Phylogenomic relationships and evolution of polyploid *Salix* species revealed by RAD sequencing data. Frontiers Plant Sciences, 11, 1077.

Whiteley, A.R., Fitzpatrick, S.W., Funk, W.C., Tallmon, D.A. (2015) Genetic rescue to the rescue. Trends in Ecology and Evolution, 30, 42–49.

Záveská, E., Maylandt, C., Paun, O., Bertel, C., Božo, F., The STEPPE Consortium, Schönswetter, P. (2019) Multiple auto- and alloploidisations marked the Pleistocene history of the widespread Eurasian steppe plant *Astragalus onobrychis* (Fabaceae). Molecular Phylogenetics and Evolution, 139, 106572.

