## Supplementary Appendix for "Little hope for the polyploid endemic Pyrenean Larkspur (*Delphinium montanum*): evidences from population genomics and Ecological Niche Modelling" (from another version of this preprint: version 1, 2021-09-16)

### Supplementary Appendix S1

Individuals of *Delphinium montanum* sampled in this study. Each row represents an individual for which sampling locality, geographic coordinates and collector name(s) are reported. There are 106 individuals in total.

| Individual ID | Locality | Longitude (°) | Latitude (°) | Elevation (m) | Collector |
|---|---|---|---|---|---|
| 20-Dmo-001 | Orri de Baix | 2.122 | 42.446 | 2170 | JB |
| 20-Dmo-002 | Orri de Baix | 2.122 | 42.446 | 2171 | JB |
| 20-Dmo-003 | Orri de Baix | 2.122 | 42.446 | 2170 | JB |
| 20-Dmo-004 | Orri de Baix | 2.122 | 42.446 | 2171 | JB |
| 20-Dmo-005 | Orri de Baix | 2.122 | 42.446 | 2175 | JB |
| 20-Dmo-006 | Orri de Baix | 2.123 | 42.446 | 2171 | JB |
| 20-Dmo-007 | Orri de Baix | 2.114 | 42.444 | 2020 | JB |
| 20-Dmo-008 | Orri de Baix | 2.114 | 42.444 | 2017 | JB |
| 20-Dmo-009 | Orri de Baix | 2.115 | 42.443 | 2036 | JB |
| 20-Dmo-010 | Orri de Baix | 2.115 | 42.443 | 2053 | JB |
| 20-Dmo-011 | Orri de Baix | 2.117 | 42.442 | 2108 | JB |
| 20-Dmo-012 | Orri de Baix | 2.117 | 42.442 | 2117 | JB |
| 20-Dmo-013 | Nohèdes | 2.263 | 42.616 | 1755 | JB |
| 20-Dmo-014 | Nohèdes | 2.263 | 42.616 | 1758 | JB |
| 20-Dmo-015 | Nohèdes | 2.263 | 42.616 | 1760 | JB |
| 20-Dmo-016 | Nohèdes | 2.263 | 42.615 | 1760 | JB |
| 20-Dmo-017 | Nohèdes | 2.263 | 42.615 | 1761 | JB |
| 20-Dmo-018 | Nohèdes | 2.263 | 42.615 | 1759 | JB |
| 20-Dmo-019 | Nohèdes | 2.263 | 42.615 | 1758 | JB |
| 20-Dmo-020 | Nohèdes | 2.263 | 42.615 | 1759 | JB |
| 20-Dmo-021 | Nohèdes | 2.263 | 42.615 | 1760 | JB |
| 20-Dmo-022 | Nohèdes | 2.263 | 42.615 | 1758 | JB |
| 20-Dmo-023 | Nohèdes | 2.263 | 42.616 | 1757 | JB |
| 20-Dmo-024 | Nohèdes | 2.263 | 42.616 | 1753 | JB |
| 20-Dmo-025 | Vallter | 2.266 | 42.427 | 2145 | AV |
| 20-Dmo-026 | Vallter | 2.266 | 42.427 | 2145 | AV |
| 20-Dmo-027 | Vallter | 2.264 | 42.426 | 2140 | AV |
| 20-Dmo-028 | Vallter | 2.264 | 42.426 | 2140 | AV |
| 20-Dmo-029 | Vallter | 2.264 | 42.426 | 2140 | AV |
| 20-Dmo-030 | Vallter | 2.264 | 42.426 | 2140 | AV |
| 20-Dmo-031 | Vallter | 2.264 | 42.426 | 2140 | AV |
| 20-Dmo-032 | Vallter | 2.264 | 42.426 | 2140 | AV |
| 20-Dmo-033 | Vallter | 2.264 | 42.426 | 2140 | AV |
| 20-Dmo-034 | Vallter | 2.264 | 42.426 | 2140 | AV |
| 20-Dmo-035 | Vallter | 2.264 | 42.426 | 2140 | AV |
| 20-Dmo-036 | Vallter | 2.264 | 42.426 | 2140 | AV |
| 20-Dmo-037 | Pedraforca | NA | NA | NA | PA, IA |
| 20-Dmo-038 | Pedraforca | 1.701 | 42.238 | NA | PA, IA |
| 20-Dmo-039 | Pedraforca | 1.705 | 42.236 | NA | PA, IA |
| 20-Dmo-040 | Serra Pedregosa | 1.688 | 42.277 | NA | PA, IA |
| 20-Dmo-041 | Cambre d'Aze | 2.131 | 42.451 | 2723 | JP, LS |
| 20-Dmo-042 | Cambre d'Aze | 2.131 | 42.451 | 2720 | JP, LS |
| 20-Dmo-043 | Cambre d'Aze | 2.124 | 42.456 | 2405 | JP, LS |
| 20-Dmo-044 | Cambre d'Aze | 2.124 | 42.456 | 2314 | JP, LS |
| 20-Dmo-045 | Serra Pedregosa | 1.688 | 42.276 | NA | PA, IA |
| 20-Dmo-046 | Tosa | 1.886 | 42.315 | NA | PA, IA |
| 20-Dmo-047 | Tosa | 1.886 | 42.315 | NA | PA, IA |
| 20-Dmo-048 | Batanist | 1.688 | 42.291 | NA | PA, IA |
| 20-Dmo-049 | Cambre d'Aze | 2.130 | 42.450 | 2649 | JP, LS |
| 20-Dmo-050 | Cambre d'Aze | 2.130 | 42.450 | 2637 | JP, LS |

| | | | | | |
|---|---|---|---|---|---|
| 20-Dmo-051 | Cambre d'Aze | 2.130 | 42.450 | 2629 | JP, LS |
| 20-Dmo-052 | Cambre d'Aze | 2.130 | 42.450 | 2626 | JP, LS |
| 20-Dmo-053 | Cambre d'Aze | 2.129 | 42.449 | 2558 | JP, LS |
| 20-Dmo-054 | Cambre d'Aze | 2.129 | 42.449 | 2585 | JP, LS |
| 20-Dmo-055 | Cambre d'Aze | 2.130 | 42.449 | 2587 | JP, LS |
| 20-Dmo-056 | Cambre d'Aze | 2.130 | 42.449 | 2598 | JP, LS |
| 20-Dmo-057 | Pedraforca | 1.706 | 42.237 | NA | JP, LS |
| 20-Dmo-058 | Pedraforca | 1.705 | 42.236 | NA | JP, LS |
| 20-Dmo-059 | Pedraforca | 1.706 | 42.236 | NA | JP, LS |
| 20-Dmo-060 | Pedraforca | 1.706 | 42.236 | NA | JP, LS |
| 20-Dmo-061 | Pedraforca | 1.703 | 42.237 | NA | JP, LS |
| 20-Dmo-062 | Pedraforca | 1.705 | 42.238 | NA | JP, LS |
| 20-Dmo-065 | Tosa | 1.886 | 42.315 | NA | PA, IA, QT |
| 20-Dmo-066 | Tosa | 1.886 | 42.315 | NA | PA, IA, QT |
| 20-Dmo-067 | Tosa | 1.886 | 42.315 | NA | PA, IA, QT |
| 20-Dmo-068 | Bastanist | 1.688 | 42.289 | NA | PA, IA |
| 20-Dmo-069 | Bastanist | 1.688 | 42.289 | NA | PA, IA |
| 20-Dmo-070 | Bastanist | 1.688 | 42.289 | NA | PA, IA |
| 20-Dmo-071 | Bastanist | 1.688 | 42.289 | NA | PA, IA |
| 20-Dmo-072 | Bastanist | 1.688 | 42.289 | NA | PA, IA |
| 20-Dmo-073 | Serra Pedregosa | 1.689 | 42.278 | NA | PA, IA |
| 20-Dmo-074 | Serra Pedregosa | 1.687 | 42.278 | NA | PA, IA |
| 20-Dmo-075 | Tosa | 1.886 | 42.315 | NA | PA, IA, QT |
| 20-Dmo-076 | Tosa | 1.886 | 42.315 | NA | PA, IA, QT |
| 20-Dmo-077 | Tosa | 1.886 | 42.315 | NA | PA, IA, QT |
| 20-Dmo-078 | Tosa | 1.886 | 42.315 | NA | PA, IA, QT |
| 20-Dmo-079 | Tosa | 1.886 | 42.315 | NA | PA, IA, QT |
| 20-Dmo-080 | Bastanist | 1.688 | 42.290 | NA | PA, IA |
| 20-Dmo-081 | Serra Pedregosa | 1.687 | 42.278 | NA | PA, IA |
| 20-Dmo-082 | Serra Pedregosa | 1.687 | 42.278 | NA | PA, IA |
| 20-Dmo-083 | Serra Pedregosa | 1.687 | 42.278 | NA | PA, IA |
| 20-Dmo-084 | Serra Pedregosa | 1.687 | 42.278 | NA | PA, IA |
| 20-Dmo-085 | Tosa | 1.886 | 42.315 | NA | PA, IA, QT |
| 20-Dmo-086 | Tosa | 1.886 | 42.315 | NA | PA, IA, QT |
| 20-Dmo-087 | Bastanist | 1.688 | 42.289 | NA | PA, IA |
| 20-Dmo-088 | Bastanist | 1.688 | 42.290 | NA | PA, IA |
| 20-Dmo-089 | Pedraforca | 1.701 | 42.238 | NA | PA, IA |
| 20-Dmo-090 | Serra Pedregosa | 1.688 | 42.277 | NA | PA, IA |
| 20-Dmo-091 | Serra Pedregosa | 1.688 | 42.277 | NA | PA, IA |
| 20-Dmo-092 | Serra Pedregosa | 1.687 | 42.277 | NA | PA, IA |
| 20-Dmo-093 | Serra Pedregosa | 1.687 | 42.277 | NA | PA, IA |
| 20-Dmo-094 | Bastanist | 1.689 | 42.288 | NA | PA, IA |
| 20-Dmo-095 | Bastanist | 1.689 | 42.288 | NA | PA, IA |
| 20-Dmo-096 | Bastanist | 1.688 | 42.291 | NA | PA, IA |
| 20-Dmo-097 | Torreta de Cadí | 1.574 | 42.282 | NA | PA, IA |
| 20-Dmo-098 | Torreta de Cadí | 1.574 | 42.283 | NA | PA, IA |
| 20-Dmo-099 | Torreta de Cadí | 1.574 | 42.283 | NA | PA, IA |
| 20-Dmo-100 | Torreta de Cadí | 1.574 | 42.282 | NA | PA, IA |
| 20-Dmo-101 | Torreta de Cadí | 1.574 | 42.283 | NA | PA, IA |
| 20-Dmo-102 | Torreta de Cadí | 1.574 | 42.282 | NA | PA, IA |
| 20-Dmo-103 | Torreta de Cadí | 1.574 | 42.282 | NA | PA, IA |
| 20-Dmo-104 | Torreta de Cadí | 1.574 | 42.282 | NA | PA, IA |
| 20-Dmo-105 | Torreta de Cadí | 1.574 | 42.283 | NA | PA, IA |
| 20-Dmo-106 | Torreta de Cadí | 1.574 | 42.283 | NA | PA, IA |
| 20-Dmo-107 | Torreta de Cadí | 1.574 | 42.283 | NA | PA, IA |
| 20-Dmo-108 | Torreta de Cadí | 1.574 | 42.282 | NA | PA, IA |

Geographic coordinates are in WGS84.

Collectors' names abbreviations: JB (Joris Bertrand), PA (Pere Aymerich), AV (Albert Vila), JP (Josep Parrera), IA (Irene Abad), Lucie Schaad and QT (Queralt Tor)

### Supplementary Appendix S2

Evolution of mean read depth depending on several combination of the key parameters of stacks (*M*, *n* and *m*) on a representative subset of 12 individuals. Individuals with minimum and maximum mean read depth are indicated in bold.

| Individual | M1n1m3 | M2n1m3 | M2n2m3 | M3n3m3 | M4n4m3 | M5n5m3 | M6n6m3 | M7n7m3 | M8n8m3 | M9n9m3 |
|---|---|---|---|---|---|---|---|---|---|---|
| 20-Dmo-001 | 27.93 | 29.42 | 29.42 | 30.51 | 31.41 | 32.25 | 32.94 | 33.47 | 33.9 | 34.25 |
| 20-Dmo-013 | 32.77 | 34.4 | 34.4 | 35.65 | 36.7 | 37.67 | 38.46 | 39.14 | 39.65 | 40.04 |
| 20-Dmo-025 | 35.86 | 37.56 | 37.55 | 38.83 | 39.84 | 40.78 | 41.55 | 41.14 | 42.61 | 43 |
| 20-Dmo-037 | 38.55 | 40.52 | 40.53 | 41.95 | 43.08 | 44.1 | 44.95 | 45.67 | 46.2 | 46.61 |
| 20-Dmo-040 | 34.5 | 36.53 | 36.53 | 37.93 | 38.97 | 39.91 | 40.74 | 41.38 | 41.9 | 42.35 |
| 20-Dmo-041 | 34.59 | 36.19 | 36.19 | 37.35 | 38.46 | 39.37 | 40.17 | 40.79 | 41.3 | 41.67 |
| 20-Dmo-046 | 30.29 | 31.84 | 31.84 | 32.96 | 33.81 | 34.56 | 35.24 | 35.77 | 36.18 | 36.55 |
| 20-Dmo-050 | 29.15 | 30.78 | 30.78 | 31.94 | 32.93 | 33.79 | 34.59 | 35.17 | 35.62 | 35.94 |
| 20-Dmo-059 | 25.36 | 26.75 | 26.75 | 27.83 | 28.73 | 29.53 | 30.21 | 30.81 | 31.25 | 31.6 |
| 20-Dmo-068 | 35.19 | 37.19 | 37.19 | 38.53 | 39.62 | 40.54 | 41.36 | 42.01 | 42.52 | 42.92 |
| 20-Dmo-097 | 26.43 | 27.78 | 27.78 | 28.79 | 29.73 | 30.54 | 31.27 | 31.84 | 32.25 | 32.63 |
| 20-Dmo-105 | 49.54 | 52.23 | 52.23 | 53.89 | 55.23 | 56.37 | 57.25 | 57.95 | 58.55 | 58.95 |

-*m*: minimum number of identical raw reads required to form a putative allele

-*M*: number of mismatches allowed between alleles to form a locus

-*n*: number of mismatches allowed between loci during construction of the catalog

Optimization of several key parameters of stacks (*M*, *n* and *m*) on a representative subset of 12 individuals. The number of total loci, removed loci, kept loci, sites filtered sites, SNPs, assembled loci, polymorphic loci and new polymorphic loci compared to the previous combination of parameters are indicated. Data were filtered with the *populations* script from STACKS following the '*80% rule*' (*i.e.* keeping only loci shared by at least 80% of samples). The selected combination is indicated in bold.

| **Parameters** | | | **Total loci** | **Removed loci** | **Kept loci** | **Sites** | **Filtered sites** | **SNPs** | **Assembled loci** | **Polymorphic loci** | **New Polymorphic loci** |
|---|---|---|---|---|---|---|---|---|---|---|---|
| **M** | **n** | **m** | | | | | | | | | |
| 1 | 1 | 3 | 91451 | 71781 | 19670 | 6692290 | 1455 | 16697 | 19669 | 10798 | NA |
| 2 | 1 | 3 | 87320 | 68155 | 19165 | 6530249 | 2555 | 19330 | 19165 | 11220 | 422 |
| 2 | 2 | 3 | 84607 | 64906 | 19701 | 6703995 | 2768 | 20496 | 19701 | 11670 | 450 |
| 3 | 3 | 3 | 81420 | 62332 | 19088 | 6498001 | 4452 | 22911 | 19088 | 11825 | 155 |
| 4 | 4 | 3 | 79107 | 60650 | 18457 | 6288499 | 6817 | 24776 | 18457 | 11825 | 0 |
| 5 | 5 | 3 | 77437 | 59628 | 17809 | 6075321 | 9073 | 26667 | 17809 | 11711 | -114 |
| 6 | 6 | 3 | 76078 | 58776 | 17302 | 5904844 | 11147 | 28488 | 17302 | 11649 | -62 |
| 7 | 7 | 3 | 74975 | 58063 | 16912 | 5778683 | 13033 | 29671 | 16912 | 11579 | -70 |
| 8 | 8 | 3 | 74289 | 57621 | 16668 | 5698383 | 14692 | 31317 | 16667 | 11563 | -16 |
| 9 | 9 | 3 | 73421 | 57029 | 16392 | 5610276 | 16184 | 32093 | 16391 | 11489 | -74 |

### Supplementary Appendix S3

Pairwise genetic differentiation ($G_{ST}$ values; Nei, 1987, $G'_{ST}$; Hendrick, 2005, $G''_{ST}$; Meirmans & Hedrick, 2011 and Jost's *D*) and pairwise geographic distances (in km). All values were found to be statistically significant ($p < 0.05$).

| $G_{ST}$/Distance | Orri de Baix | Nohèdes | Vallter | Pedraforca | Serra Pedregosa | Cambre d'Aze | Tosa | Bastanist | Torreta de Cadi |
|---|---|---|---|---|---|---|---|---|---|
| Orri de Baix | | 0.441 | 0.433 | 0.394 | 0.390 | 0.156 | 0.427 | 0.382 | 0.483 |
| Nohèdes | 22 | | 0.480 | 0.439 | 0.437 | 0.404 | 0.493 | 0.436 | 0.527 |
| Vallter | 12 | 21 | | 0.426 | 0.438 | 0.401 | 0.490 | 0.426 | 0.518 |
| Pedraforca | 41 | 62 | 51 | | 0.264 | 0.357 | 0.383 | 0.264 | 0.360 |
| Serra Pedregosa | 40 | 60 | 50 | 5 | | 0.360 | 0.380 | 0.153 | 0.376 |
| Cambre d'Aze | 1 | 21 | 11 | 42 | 41 | | 0.394 | 0.353 | 0.449 |
| Tosa | 24 | 45 | 33 | 17 | 17 | 25 | | 0.375 | 0.480 |
| Bastanist | 39 | 59 | 50 | 6 | 1 | 40 | 17 | | 0.369 |
| Torreta de Cadi | 48 | 68 | 59 | 12 | 9 | 49 | 26 | 9 | |

| $G'_{ST}$/Distance | Orri de Baix | Nohèdes | Vallter | Pedraforca | Serra Pedregosa | Cambre d'Aze | Tosa | Bastanist | Torreta de Cadi |
|---|---|---|---|---|---|---|---|---|---|
| Orri de Baix | | 0.452 | 0.443 | 0.404 | 0.400 | 0.172 | 0.436 | 0.393 | 0.493 |
| Nohèdes | 22 | | 0.490 | 0.447 | 0.448 | 0.416 | 0.503 | 0.447 | 0.537 |
| Vallter | 12 | 21 | | 0.434 | 0.448 | 0.412 | 0.499 | 0.437 | 0.527 |
| Pedraforca | 41 | 62 | 51 | | 0.280 | 0.371 | 0.393 | 0.279 | 0.372 |
| Serra Pedregosa | 40 | 60 | 50 | 5 | | 0.372 | 0.391 | 0.167 | 0.387 |
| Cambre d'Aze | 1 | 21 | 11 | 42 | 41 | | 0.405 | 0.365 | 0.46 |
| Tosa | 24 | 45 | 33 | 17 | 17 | 25 | | 0.386 | 0.491 |
| Bastanist | 39 | 59 | 50 | 6 | 1 | 40 | 17 | | 0.38 |

| | | | | | | | | |
|---|---|---|---|---|---|---|---|---|
| Torreta de Cadi | 48 | 68 | 59 | 12 | 9 | 49 | 26 | 9 |

| $G''_{ST}$/ Jost's *D* | Orri de Baix | Nohèdes | Vallter | Pedraforca | Serra Pedregosa | Cambre d'Aze | Tosa | Bastanist | Torreta de Cadi |
|---|---|---|---|---|---|---|---|---|---|
| Orri de Baix | | 0.542 | 0.531 | 0.514 | 0.507 | 0.214 | 0.533 | 0.499 | 0.606 |
| Nohèdes | 0.163 | | 0.579 | 0.559 | 0.559 | 0.510 | 0.604 | 0.559 | 0.650 |
| Vallter | 0.158 | 0.172 | | 0.543 | 0.559 | 0.505 | 0.600 | 0.546 | 0.639 |
| Pedraforca | 0.185 | 0.202 | 0.193 | | 0.372 | 0.484 | 0.501 | 0.372 | 0.479 |
| Serra Pedregosa | 0.178 | 0.199 | 0.200 | 0.127 | | 0.483 | 0.498 | 0.221 | 0.496 |
| Cambre d'Aze | 0.051 | 0.159 | 0.158 | 0.180 | 0.177 | | 0.506 | 0.475 | 0.579 |
| Tosa | 0.171 | 0.202 | 0.201 | 0.179 | 0.174 | 0.169 | | 0.492 | 0.605 |
| Bastanist | 0.175 | 0.201 | 0.194 | 0.128 | 0.065 | 0.174 | 0.172 | | 0.489 |
| Torreta de Cadi | 0.223 | 0.243 | 0.235 | 0.169 | 0.176 | 0.220 | 0.224 | 0.174 | |

### Supplementary Appendix S4

Summary of the 135 models of Maxent ENMs generated by ENMeval. The two best models are indicated in bold.

| | settings | features | rm | train.AUC | avg.test.AUC | var.test.AUC | avg.diff.AUC | var.diff.AUC | avg.test.orMTP | var.test.orMTP | avg.test.or10pct | var.test.or10pct | AICc | delta.AICc | w.AIC | parameters | sel.cri | rankAIC |
|---|---|---|---|---|---|---|---|---|---|---|---|---|---|---|---|---|---|---|
| **1** | L_0.5 | L | 0.5 | 0.992 | 0.984 | 0.000 | 0.008 | 0.001 | 0.250 | 0.250 | 0.375 | 0.229 | 164.270 | 12.016 | 0.001 | 4 | | 18 |
| **2** | P_0.5 | P | 0.5 | 0.992 | 0.984 | 0.000 | 0.006 | 0.000 | 0.250 | 0.083 | 0.375 | 0.229 | 179.791 | 27.537 | 0.000 | 5 | | 96 |
| **3** | Q_0.5 | Q | 0.5 | 0.993 | 0.986 | 0.000 | 0.007 | 0.000 | 0.250 | 0.250 | 0.375 | 0.229 | 161.481 | 9.228 | 0.003 | 4 | | 14 |
| **4** | H_0.5 | H | 0.5 | 0.997 | 0.988 | 0.000 | 0.006 | 0.000 | 0.250 | 0.250 | 0.375 | 0.229 | NA | NA | NA | 9 | | 127 |
| **5** | LP_0.5 | LP | 0.5 | 0.993 | 0.984 | 0.000 | 0.007 | 0.000 | 0.250 | 0.250 | 0.375 | 0.229 | 175.714 | 23.461 | 0.000 | 5 | | 77 |
| **6** | LQ_0.5 | LQ | 0.5 | 0.994 | 0.988 | 0.000 | 0.006 | 0.000 | 0.250 | 0.250 | 0.375 | 0.229 | 158.292 | 6.038 | 0.016 | 4 | | 7 |
| **7** | LH_0.5 | LH | 0.5 | 0.997 | 0.986 | 0.000 | 0.006 | 0.000 | 0.250 | 0.250 | 0.375 | 0.229 | NA | NA | NA | 9 | | 128 |
| **8** | PQ_0.5 | PQ | 0.5 | 0.994 | 0.988 | 0.000 | 0.006 | 0.000 | 0.250 | 0.250 | 0.375 | 0.229 | 158.834 | 6.580 | 0.012 | 4 | | 9 |
| **9** | PH_0.5 | PH | 0.5 | 0.997 | 0.988 | 0.000 | 0.006 | 0.000 | 0.250 | 0.250 | 0.375 | 0.229 | NA | NA | NA | 10 | | 129 |
| **10** | QH_0.5 | QH | 0.5 | 0.997 | 0.987 | 0.000 | 0.006 | 0.000 | 0.250 | 0.250 | 0.375 | 0.229 | NA | NA | NA | 8 | | 130 |
| **11** | LPQ_0.5 | LPQ | 0.5 | 0.994 | 0.987 | 0.000 | 0.006 | 0.000 | 0.250 | 0.250 | 0.375 | 0.229 | 158.834 | 6.581 | 0.012 | 4 | | 10 |
| **12** | LPH_0.5 | LPH | 0.5 | 0.997 | 0.988 | 0.000 | 0.006 | 0.000 | 0.250 | 0.250 | 0.375 | 0.229 | NA | NA | NA | 10 | | 131 |
| **13** | LQH_0.5 | LQH | 0.5 | 0.997 | 0.987 | 0.000 | 0.006 | 0.000 | 0.250 | 0.250 | 0.375 | 0.229 | NA | NA | NA | 8 | | 132 |
| **14** | PQH_0.5 | PQH | 0.5 | 0.997 | 0.988 | 0.000 | 0.007 | 0.000 | 0.250 | 0.250 | 0.375 | 0.229 | NA | NA | NA | 9 | | 133 |
| **15** | LPQH_0.5 | LPQH | 0.5 | 0.997 | 0.988 | 0.000 | 0.007 | 0.000 | 0.250 | 0.250 | 0.375 | 0.229 | NA | NA | NA | 10 | | 134 |
| **16** | L_1 | L | 1 | 0.992 | 0.982 | 0.000 | 0.008 | 0.000 | 0.250 | 0.250 | 0.375 | 0.229 | 167.297 | 15.043 | 0.000 | 4 | | 32 |
| **17** | P_1 | P | 1 | 0.989 | 0.979 | 0.000 | 0.007 | 0.000 | 0.250 | 0.083 | 0.333 | 0.056 | 190.123 | 37.869 | 0.000 | 5 | | 110 |
| **18** | Q_1 | Q | 1 | 0.993 | 0.985 | 0.000 | 0.007 | 0.000 | 0.375 | 0.229 | 0.375 | 0.229 | 163.697 | 11.443 | 0.001 | 4 | | 17 |
| **19** | H_1 | H | 1 | 0.996 | 0.987 | 0.000 | 0.005 | 0.000 | 0.250 | 0.250 | 0.250 | 0.250 | 262.110 | 109.857 | 0.000 | 7 | | 125 |
| **20** | LP_1 | LP | 1 | 0.990 | 0.980 | 0.000 | 0.008 | 0.000 | 0.375 | 0.229 | 0.542 | 0.174 | 186.641 | 34.388 | 0.000 | 5 | | 106 |
| **21** | LQ_1 | LQ | 1 | 0.994 | 0.987 | 0.000 | 0.006 | 0.000 | 0.250 | 0.250 | 0.375 | 0.229 | 160.280 | 8.026 | 0.006 | 4 | | 11 |
| **22** | LH_1 | LH | 1 | 0.995 | 0.985 | 0.000 | 0.006 | 0.000 | 0.250 | 0.250 | 0.250 | 0.250 | NA | NA | NA | 8 | | 135 |
| **23** | PQ_1 | PQ | 1 | 0.993 | 0.980 | 0.000 | 0.008 | 0.000 | 0.375 | 0.229 | 0.542 | 0.174 | 166.411 | 14.157 | 0.000 | 4 | | 28 |

| | | | | | | | | | | | | | | | | | | |
|---|---|---|---|---|---|---|---|---|---|---|---|---|---|---|---|---|---|---|
| **24** | PH_1 | PH | | 1 | 0.995 | 0.988 | 0.000 | 0.006 | 0.000 | 0.250 | 0.250 | 0.375 | 0.229 | 262.704 | 110.451 | 0.000 | 7 | 126 |
| **25** | QH_1 | QH | | 1 | 0.995 | 0.986 | 0.000 | 0.005 | 0.000 | 0.250 | 0.250 | 0.375 | 0.229 | 165.298 | 13.044 | 0.000 | 5 | 27 |
| **26** | LPQ_1 | LPQ | | 1 | 0.993 | 0.980 | 0.000 | 0.008 | 0.000 | 0.375 | 0.229 | 0.542 | 0.174 | 166.411 | 14.157 | 0.000 | 4 | 29 |
| **27** | LPH_1 | LPH | | 1 | 0.995 | 0.987 | 0.000 | 0.006 | 0.000 | 0.250 | 0.250 | 0.375 | 0.229 | 190.731 | 38.477 | 0.000 | 6 | 114 |
| **28** | LQH_1 | LQH | | 1 | 0.995 | 0.986 | 0.000 | 0.005 | 0.000 | 0.250 | 0.250 | 0.375 | 0.229 | 165.294 | 13.041 | 0.000 | 5 | 26 |
| **29** | PQH_1 | PQH | | 1 | 0.995 | 0.987 | 0.000 | 0.006 | 0.000 | 0.250 | 0.250 | 0.375 | 0.229 | 190.594 | 38.341 | 0.000 | 6 | 111 |
| **30** | LPQH_1 | LPQH | | 1 | 0.995 | 0.987 | 0.000 | 0.006 | 0.000 | 0.250 | 0.250 | 0.375 | 0.229 | 190.594 | 38.341 | 0.000 | 6 | 112 |
| **31** | L_1.5 | L | 1.5 | | 0.991 | 0.978 | 0.000 | 0.009 | 0.000 | 0.375 | 0.229 | 0.625 | 0.229 | 171.977 | 19.724 | 0.000 | 4 | 55 |
| **32** | P_1.5 | P | 1.5 | | 0.982 | 0.978 | 0.000 | 0.008 | 0.000 | 0.250 | 0.083 | 0.417 | 0.083 | 180.445 | 28.191 | 0.000 | 3 | 99 |
| **33** | Q_1.5 | Q | 1.5 | | 0.992 | 0.981 | 0.000 | 0.008 | 0.000 | 0.375 | 0.229 | 0.542 | 0.174 | 167.003 | 14.749 | 0.000 | 4 | 30 |
| **34** | H_1.5 | H | 1.5 | | 0.995 | 0.986 | 0.000 | 0.006 | 0.000 | 0.250 | 0.250 | 0.250 | 0.250 | 195.376 | 43.123 | 0.000 | 6 | 122 |
| **35** | LP_1.5 | LP | 1.5 | | 0.984 | 0.978 | 0.000 | 0.008 | 0.000 | 0.250 | 0.083 | 0.417 | 0.083 | 174.697 | 22.443 | 0.000 | 3 | 72 |
| **36** | LQ_1.5 | LQ | 1.5 | | 0.994 | 0.984 | 0.000 | 0.007 | 0.000 | 0.375 | 0.229 | 0.375 | 0.229 | 163.206 | 10.952 | 0.001 | 4 | 15 |
| **37** | LH_1.5 | LH | 1.5 | | 0.995 | 0.984 | 0.000 | 0.007 | 0.000 | 0.250 | 0.250 | 0.375 | 0.229 | 194.601 | 42.347 | 0.000 | 6 | 121 |
| **38** | PQ_1.5 | PQ | 1.5 | | 0.986 | 0.979 | 0.000 | 0.009 | 0.000 | 0.375 | 0.229 | 0.625 | 0.229 | 178.614 | 26.360 | 0.000 | 4 | 92 |
| **39** | PH_1.5 | PH | 1.5 | | 0.995 | 0.988 | 0.000 | 0.005 | 0.000 | 0.250 | 0.250 | 0.375 | 0.229 | 158.595 | 6.341 | 0.014 | 4 | 8 |
| **40** | QH_1.5 | QH | 1.5 | | 0.995 | 0.987 | 0.000 | 0.005 | 0.000 | 0.250 | 0.250 | 0.375 | 0.229 | 193.233 | 40.980 | 0.000 | 6 | 118 |
| **41** | LPQ_1.5 | LPQ | 1.5 | | 0.986 | 0.979 | 0.000 | 0.009 | 0.000 | 0.375 | 0.229 | 0.542 | 0.174 | 178.614 | 26.360 | 0.000 | 4 | 93 |
| **42** | LPH_1.5 | LPH | 1.5 | | 0.995 | 0.986 | 0.000 | 0.006 | 0.000 | 0.250 | 0.250 | 0.375 | 0.229 | 171.530 | 19.276 | 0.000 | 5 | 52 |
| **43** | LQH_1.5 | LQH | 1.5 | | 0.995 | 0.987 | 0.000 | 0.005 | 0.000 | 0.250 | 0.250 | 0.375 | 0.229 | 193.233 | 40.980 | 0.000 | 6 | 119 |
| **44** | PQH_1.5 | PQH | 1.5 | | 0.994 | 0.986 | 0.000 | 0.006 | 0.000 | 0.250 | 0.250 | 0.500 | 0.167 | 195.671 | 43.417 | 0.000 | 6 | 123 |
| **45** | LPQH_1.5 | LPQH | 1.5 | | 0.994 | 0.985 | 0.000 | 0.006 | 0.000 | 0.250 | 0.250 | 0.500 | 0.167 | 195.671 | 43.417 | 0.000 | 6 | 124 |
| **46** | L_2 | L | | 2 | 0.987 | 0.976 | 0.000 | 0.010 | 0.000 | 0.375 | 0.229 | 0.625 | 0.229 | 178.226 | 25.972 | 0.000 | 4 | 91 |
| **47** | P_2 | P | | 2 | 0.982 | 0.977 | 0.000 | 0.009 | 0.000 | 0.125 | 0.063 | 0.500 | 0.167 | 181.826 | 29.572 | 0.000 | 3 | 100 |
| **48** | Q_2 | Q | | 2 | 0.992 | 0.980 | 0.000 | 0.009 | 0.000 | 0.375 | 0.229 | 0.625 | 0.229 | 171.343 | 19.089 | 0.000 | 4 | 51 |
| **49** | H_2 | H | | 2 | 0.994 | 0.982 | 0.000 | 0.008 | 0.000 | 0.250 | 0.250 | 0.375 | 0.229 | 177.171 | 24.917 | 0.000 | 5 | 85 |
| **50** | LP_2 | LP | | 2 | 0.982 | 0.978 | 0.000 | 0.007 | 0.000 | 0.250 | 0.083 | 0.500 | 0.167 | 172.358 | 20.105 | 0.000 | 2 | 59 |
| **51** | LQ_2 | LQ | | 2 | 0.993 | 0.981 | 0.000 | 0.008 | 0.000 | 0.375 | 0.229 | 0.625 | 0.229 | 167.187 | 14.933 | 0.000 | 4 | 31 |

| | | | | | | | | | | | | | | | | | | | |
|---|---|---|---|---|---|---|---|---|---|---|---|---|---|---|---|---|---|---|---|
| **52** | LH_2 | LH | | 2 | 0.994 | 0.984 | 0.000 | 0.007 | 0.000 | 0.250 | 0.250 | 0.375 | 0.229 | 163.386 | 11.132 | 0.001 | 4 | | 16 |
| **53** | PQ_2 | PQ | | 2 | 0.983 | 0.979 | 0.000 | 0.007 | 0.000 | 0.250 | 0.083 | 0.500 | 0.167 | 170.000 | 17.746 | 0.000 | 2 | | 46 |
| **54** | **PH_2** | **PH** | | **2** | **0.995** | **0.989** | **0.000** | **0.005** | **0.000** | **0.250** | **0.250** | **0.375** | **0.229** | **174.628** | **22.374** | **0.000** | **5** | **AUCmtp. AUC10** | **71** |
| **55** | **QH_2** | **QH** | | **2** | **0.994** | **0.988** | **0.000** | **0.005** | **0.000** | **0.250** | **0.250** | **0.375** | **0.229** | **152.254** | **0.000** | **0.327** | **3** | **LowAIC** | **1** |
| **56** | LPQ_2 | LPQ | | 2 | 0.983 | 0.979 | 0.000 | 0.008 | 0.000 | 0.250 | 0.083 | 0.500 | 0.167 | 170.000 | 17.746 | 0.000 | 2 | | 45 |
| **57** | LPH_2 | LPH | | 2 | 0.994 | 0.984 | 0.000 | 0.008 | 0.000 | 0.125 | 0.063 | 0.375 | 0.229 | 176.919 | 24.665 | 0.000 | 5 | | 84 |
| **58** | LQH_2 | LQH | | 2 | 0.994 | 0.988 | 0.000 | 0.005 | 0.000 | 0.250 | 0.250 | 0.375 | 0.229 | 152.254 | 0.000 | 0.327 | 3 | | 2 |
| **59** | PQH_2 | PQH | | 2 | 0.993 | 0.985 | 0.000 | 0.006 | 0.000 | 0.125 | 0.063 | 0.375 | 0.229 | 164.505 | 12.252 | 0.001 | 4 | | 21 |
| **60** | LPQH_2 | LPQH | | 2 | 0.993 | 0.985 | 0.000 | 0.007 | 0.000 | 0.250 | 0.250 | 0.375 | 0.229 | 164.505 | 12.252 | 0.001 | 4 | | 22 |
| **61** | L_2.5 | L | 2.5 | | 0.981 | 0.973 | 0.000 | 0.011 | 0.000 | 0.375 | 0.229 | 0.625 | 0.229 | 172.343 | 20.089 | 0.000 | 2 | | 58 |
| **62** | P_2.5 | P | 2.5 | | 0.982 | 0.974 | 0.000 | 0.009 | 0.001 | 0.125 | 0.063 | 0.375 | 0.229 | 183.538 | 31.284 | 0.000 | 3 | | 101 |
| **63** | Q_2.5 | Q | 2.5 | | 0.989 | 0.978 | 0.000 | 0.009 | 0.000 | 0.375 | 0.229 | 0.625 | 0.229 | 168.380 | 16.127 | 0.000 | 3 | | 36 |
| **64** | H_2.5 | H | 2.5 | | 0.993 | 0.975 | 0.000 | 0.011 | 0.000 | 0.250 | 0.250 | 0.625 | 0.229 | 172.408 | 20.154 | 0.000 | 4 | | 60 |
| **65** | LP_2.5 | LP | 2.5 | | 0.982 | 0.978 | 0.000 | 0.006 | 0.000 | 0.250 | 0.083 | 0.500 | 0.167 | 173.475 | 21.222 | 0.000 | 2 | | 66 |
| **66** | LQ_2.5 | LQ | 2.5 | | 0.992 | 0.978 | 0.000 | 0.010 | 0.000 | 0.375 | 0.229 | 0.625 | 0.229 | 164.945 | 12.691 | 0.001 | 3 | | 25 |
| **67** | LH_2.5 | LH | 2.5 | | 0.993 | 0.980 | 0.000 | 0.009 | 0.000 | 0.250 | 0.250 | 0.375 | 0.229 | 179.937 | 27.683 | 0.000 | 5 | | 98 |
| **68** | PQ_2.5 | PQ | 2.5 | | 0.982 | 0.978 | 0.000 | 0.006 | 0.000 | 0.125 | 0.063 | 0.500 | 0.167 | 171.077 | 18.823 | 0.000 | 2 | | 49 |
| **69** | PH_2.5 | PH | 2.5 | | 0.995 | 0.985 | 0.000 | 0.007 | 0.000 | 0.250 | 0.250 | 0.458 | 0.174 | 179.824 | 27.571 | 0.000 | 5 | | 97 |
| **70** | QH_2.5 | QH | 2.5 | | 0.994 | 0.987 | 0.000 | 0.005 | 0.000 | 0.250 | 0.250 | 0.375 | 0.229 | 154.566 | 2.313 | 0.103 | 3 | | 3 |
| **71** | LPQ_2.5 | LPQ | 2.5 | | 0.982 | 0.978 | 0.000 | 0.007 | 0.000 | 0.250 | 0.083 | 0.500 | 0.167 | 171.077 | 18.823 | 0.000 | 2 | | 50 |
| **72** | LPH_2.5 | LPH | 2.5 | | 0.992 | 0.980 | 0.000 | 0.009 | 0.000 | 0.125 | 0.063 | 0.542 | 0.174 | 171.909 | 19.655 | 0.000 | 4 | | 54 |
| **73** | LQH_2.5 | LQH | 2.5 | | 0.994 | 0.987 | 0.000 | 0.005 | 0.000 | 0.250 | 0.250 | 0.375 | 0.229 | 154.566 | 2.313 | 0.103 | 3 | | 4 |
| **74** | PQH_2.5 | PQH | 2.5 | | 0.993 | 0.982 | 0.000 | 0.008 | 0.000 | 0.125 | 0.063 | 0.375 | 0.229 | 168.297 | 16.043 | 0.000 | 4 | | 34 |
| **75** | LPQH_2.5 | LPQH | 2.5 | | 0.993 | 0.982 | 0.000 | 0.008 | 0.000 | 0.125 | 0.063 | 0.375 | 0.229 | 168.297 | 16.043 | 0.000 | 4 | | 35 |
| **76** | L_3 | L | | 3 | 0.978 | 0.972 | 0.000 | 0.010 | 0.000 | 0.250 | 0.083 | 0.625 | 0.229 | 174.761 | 22.507 | 0.000 | 2 | | 74 |
| **77** | P_3 | P | | 3 | 0.981 | 0.972 | 0.001 | 0.010 | 0.001 | 0.125 | 0.063 | 0.375 | 0.229 | 185.564 | 33.310 | 0.000 | 3 | | 104 |
| **78** | Q_3 | Q | | 3 | 0.986 | 0.976 | 0.000 | 0.009 | 0.000 | 0.375 | 0.229 | 0.625 | 0.229 | 167.555 | 15.301 | 0.000 | 2 | | 33 |

| | | | | | | | | | | | | | | | | | | |
|---|---|---|---|---|---|---|---|---|---|---|---|---|---|---|---|---|---|---|
| **79** | H_3 | H | | 3 | 0.990 | 0.973 | 0.000 | 0.011 | 0.000 | 0.250 | 0.083 | 0.625 | 0.229 | 173.426 | 21.173 | 0.000 | 3 | 65 |
| **80** | LP_3 | LP | | 3 | 0.981 | 0.978 | 0.000 | 0.004 | 0.000 | 0.125 | 0.063 | 0.500 | 0.167 | 174.759 | 22.506 | 0.000 | 2 | 73 |
| **81** | LQ_3 | LQ | | 3 | 0.988 | 0.976 | 0.000 | 0.010 | 0.000 | 0.375 | 0.229 | 0.625 | 0.229 | 169.816 | 17.562 | 0.000 | 3 | 44 |
| **82** | LH_3 | LH | | 3 | 0.991 | 0.976 | 0.000 | 0.010 | 0.000 | 0.125 | 0.063 | 0.625 | 0.229 | 173.834 | 21.580 | 0.000 | 4 | 69 |
| **83** | PQ_3 | PQ | | 3 | 0.982 | 0.978 | 0.000 | 0.004 | 0.000 | 0.125 | 0.063 | 0.500 | 0.167 | 172.308 | 20.054 | 0.000 | 2 | 56 |
| **84** | PH_3 | PH | | 3 | 0.994 | 0.981 | 0.000 | 0.009 | 0.000 | 0.375 | 0.229 | 0.625 | 0.229 | 185.433 | 33.179 | 0.000 | 5 | 103 |
| **85** | QH_3 | QH | | 3 | 0.994 | 0.982 | 0.000 | 0.007 | 0.000 | 0.250 | 0.250 | 0.375 | 0.229 | 157.298 | 5.044 | 0.026 | 3 | 5 |
| **86** | LPQ_3 | LPQ | | 3 | 0.982 | 0.977 | 0.000 | 0.005 | 0.000 | 0.125 | 0.063 | 0.500 | 0.167 | 172.308 | 20.054 | 0.000 | 2 | 57 |
| **87** | LPH_3 | LPH | | 3 | 0.991 | 0.978 | 0.000 | 0.009 | 0.000 | 0.250 | 0.083 | 0.625 | 0.229 | 169.434 | 17.181 | 0.000 | 3 | 43 |
| **88** | LQH_3 | LQH | | 3 | 0.994 | 0.982 | 0.000 | 0.007 | 0.000 | 0.250 | 0.250 | 0.375 | 0.229 | 157.298 | 5.044 | 0.026 | 3 | 6 |
| **89** | PQH_3 | PQH | | 3 | 0.992 | 0.978 | 0.000 | 0.009 | 0.000 | 0.125 | 0.063 | 0.625 | 0.229 | 164.851 | 12.597 | 0.001 | 3 | 23 |
| **90** | LPQH_3 | LPQH | | 3 | 0.992 | 0.978 | 0.000 | 0.009 | 0.000 | 0.125 | 0.063 | 0.625 | 0.229 | 164.851 | 12.597 | 0.001 | 3 | 24 |
| **91** | L_3.5 | L | 3.5 | | 0.975 | 0.971 | 0.000 | 0.008 | 0.000 | 0.250 | 0.083 | 0.500 | 0.167 | 177.536 | 25.282 | 0.000 | 2 | 86 |
| **92** | P_3.5 | P | 3.5 | | 0.981 | 0.972 | 0.001 | 0.011 | 0.001 | 0.125 | 0.063 | 0.375 | 0.229 | 187.963 | 35.709 | 0.000 | 3 | 108 |
| **93** | Q_3.5 | Q | 3.5 | | 0.984 | 0.975 | 0.000 | 0.008 | 0.000 | 0.250 | 0.083 | 0.625 | 0.229 | 169.091 | 16.838 | 0.000 | 2 | 41 |
| **94** | H_3.5 | H | 3.5 | | 0.984 | 0.972 | 0.000 | 0.010 | 0.000 | 0.250 | 0.083 | 0.500 | 0.167 | 179.241 | 26.987 | 0.000 | 3 | 94 |
| **95** | LP_3.5 | LP | 3.5 | | 0.980 | 0.976 | 0.000 | 0.004 | 0.000 | 0.125 | 0.063 | 0.375 | 0.229 | 176.181 | 23.928 | 0.000 | 2 | 79 |
| **96** | LQ_3.5 | LQ | 3.5 | | 0.984 | 0.974 | 0.000 | 0.010 | 0.000 | 0.375 | 0.229 | 0.625 | 0.229 | 169.091 | 16.838 | 0.000 | 2 | 42 |
| **97** | LH_3.5 | LH | 3.5 | | 0.988 | 0.970 | 0.000 | 0.012 | 0.000 | 0.250 | 0.083 | 0.625 | 0.229 | 171.776 | 19.523 | 0.000 | 3 | 53 |
| **98** | PQ_3.5 | PQ | 3.5 | | 0.980 | 0.977 | 0.000 | 0.004 | 0.000 | 0.125 | 0.063 | 0.375 | 0.229 | 173.677 | 21.424 | 0.000 | 2 | 68 |
| **99** | PH_3.5 | PH | 3.5 | | 0.992 | 0.978 | 0.000 | 0.009 | 0.000 | 0.250 | 0.083 | 0.500 | 0.167 | 192.068 | 39.815 | 0.000 | 5 | 116 |
| **100** | QH_3.5 | QH | 3.5 | | 0.994 | 0.979 | 0.000 | 0.008 | 0.000 | 0.125 | 0.063 | 0.625 | 0.229 | 160.494 | 8.240 | 0.005 | 3 | 12 |
| **101** | LPQ_3.5 | LPQ | 3.5 | | 0.980 | 0.977 | 0.000 | 0.004 | 0.000 | 0.125 | 0.063 | 0.375 | 0.229 | 173.662 | 21.409 | 0.000 | 2 | 67 |
| **102** | LPH_3.5 | LPH | 3.5 | | 0.989 | 0.976 | 0.000 | 0.009 | 0.000 | 0.250 | 0.083 | 0.500 | 0.167 | 173.840 | 21.587 | 0.000 | 3 | 70 |
| **103** | LQH_3.5 | LQH | 3.5 | | 0.994 | 0.979 | 0.000 | 0.008 | 0.000 | 0.250 | 0.250 | 0.625 | 0.229 | 160.494 | 8.240 | 0.005 | 3 | 13 |
| **104** | PQH_3.5 | PQH | 3.5 | | 0.991 | 0.977 | 0.000 | 0.009 | 0.000 | 0.250 | 0.083 | 0.625 | 0.229 | 168.744 | 16.490 | 0.000 | 3 | 40 |
| **105** | LPQH_3.5 | LPQH | 3.5 | | 0.991 | 0.977 | 0.000 | 0.009 | 0.000 | 0.250 | 0.083 | 0.625 | 0.229 | 168.731 | 16.477 | 0.000 | 3 | 39 |
| **106** | L_4 | L | | 4 | 0.972 | 0.971 | 0.000 | 0.006 | 0.000 | 0.125 | 0.063 | 0.500 | 0.167 | 176.564 | 24.311 | 0.000 | 1 | 80 |

| | | | | | | | | | | | | | | | | | | |
|---|---|---|---|---|---|---|---|---|---|---|---|---|---|---|---|---|---|---|
| **107** | P_4 | P | | 4 | 0.980 | 0.971 | 0.001 | 0.012 | 0.001 | 0.125 | 0.063 | 0.375 | 0.229 | 190.705 | 38.452 | 0.000 | 3 | 113 |
| **108** | Q_4 | Q | | 4 | 0.982 | 0.973 | 0.000 | 0.007 | 0.000 | 0.125 | 0.063 | 0.500 | 0.167 | 170.783 | 18.530 | 0.000 | 2 | 48 |
| **109** | H_4 | H | | 4 | 0.976 | 0.973 | 0.000 | 0.006 | 0.000 | 0.125 | 0.063 | 0.500 | 0.167 | 192.826 | 40.572 | 0.000 | 4 | 117 |
| **110** | LP_4 | LP | | 4 | 0.979 | 0.975 | 0.000 | 0.005 | 0.000 | 0.125 | 0.063 | 0.375 | 0.229 | 177.713 | 25.460 | 0.000 | 2 | 88 |
| **111** | LQ_4 | LQ | | 4 | 0.982 | 0.972 | 0.000 | 0.010 | 0.000 | 0.250 | 0.083 | 0.625 | 0.229 | 170.783 | 18.530 | 0.000 | 2 | 47 |
| **112** | LH_4 | LH | | 4 | 0.983 | 0.971 | 0.000 | 0.011 | 0.000 | 0.250 | 0.083 | 0.625 | 0.229 | 177.574 | 25.320 | 0.000 | 3 | 87 |
| **113** | PQ_4 | PQ | | 4 | 0.979 | 0.975 | 0.000 | 0.005 | 0.000 | 0.125 | 0.063 | 0.375 | 0.229 | 175.108 | 22.854 | 0.000 | 2 | 75 |
| **114** | PH_4 | PH | | 4 | 0.986 | 0.977 | 0.000 | 0.006 | 0.000 | 0.125 | 0.063 | 0.500 | 0.167 | 186.698 | 34.444 | 0.000 | 4 | 107 |
| **115** | QH_4 | QH | | 4 | 0.993 | 0.974 | 0.000 | 0.010 | 0.000 | 0.250 | 0.083 | 0.625 | 0.229 | 164.473 | 12.219 | 0.001 | 3 | 19 |
| **116** | LPQ_4 | LPQ | | 4 | 0.979 | 0.975 | 0.000 | 0.005 | 0.000 | 0.125 | 0.063 | 0.375 | 0.229 | 175.108 | 22.854 | 0.000 | 2 | 76 |
| **117** | LPH_4 | LPH | | 4 | 0.984 | 0.974 | 0.000 | 0.009 | 0.000 | 0.250 | 0.083 | 0.500 | 0.167 | 185.913 | 33.660 | 0.000 | 4 | 105 |
| **118** | LQH_4 | LQH | | 4 | 0.993 | 0.973 | 0.000 | 0.011 | 0.000 | 0.250 | 0.083 | 0.625 | 0.229 | 164.473 | 12.219 | 0.001 | 3 | 20 |
| **119** | PQH_4 | PQH | | 4 | 0.989 | 0.974 | 0.000 | 0.010 | 0.000 | 0.250 | 0.083 | 0.500 | 0.167 | 173.046 | 20.792 | 0.000 | 3 | 63 |
| **120** | LPQH_4 | LPQH | | 4 | 0.989 | 0.974 | 0.000 | 0.010 | 0.000 | 0.250 | 0.083 | 0.500 | 0.167 | 173.046 | 20.792 | 0.000 | 3 | 64 |
| **121** | L_4.5 | L | 4.5 | | 0.972 | 0.971 | 0.000 | 0.006 | 0.000 | 0.125 | 0.063 | 0.500 | 0.167 | 176.854 | 24.600 | 0.000 | 1 | 83 |
| **122** | P_4.5 | P | 4.5 | | 0.980 | 0.969 | 0.001 | 0.012 | 0.001 | 0.125 | 0.063 | 0.375 | 0.229 | 193.856 | 41.602 | 0.000 | 3 | 120 |
| **123** | Q_4.5 | Q | 4.5 | | 0.979 | 0.972 | 0.000 | 0.006 | 0.000 | 0.125 | 0.063 | 0.500 | 0.167 | 172.617 | 20.363 | 0.000 | 2 | 61 |
| **124** | H_4.5 | H | 4.5 | | 0.972 | 0.973 | 0.000 | 0.005 | 0.000 | 0.125 | 0.063 | 0.375 | 0.229 | 188.578 | 36.324 | 0.000 | 3 | 109 |
| **125** | LP_4.5 | LP | 4.5 | | 0.977 | 0.973 | 0.000 | 0.005 | 0.000 | 0.125 | 0.063 | 0.375 | 0.229 | 179.329 | 27.075 | 0.000 | 2 | 95 |
| **126** | LQ_4.5 | LQ | 4.5 | | 0.979 | 0.972 | 0.000 | 0.007 | 0.000 | 0.125 | 0.063 | 0.500 | 0.167 | 172.617 | 20.363 | 0.000 | 2 | 62 |
| **127** | LH_4.5 | LH | 4.5 | | 0.980 | 0.970 | 0.000 | 0.010 | 0.000 | 0.250 | 0.083 | 0.500 | 0.167 | 175.906 | 23.652 | 0.000 | 2 | 78 |
| **128** | PQ_4.5 | PQ | 4.5 | | 0.978 | 0.974 | 0.000 | 0.005 | 0.000 | 0.125 | 0.063 | 0.375 | 0.229 | 176.614 | 24.360 | 0.000 | 2 | 82 |
| **129** | PH_4.5 | PH | 4.5 | | 0.979 | 0.975 | 0.000 | 0.005 | 0.000 | 0.125 | 0.063 | 0.375 | 0.229 | 192.051 | 39.797 | 0.000 | 4 | 115 |
| **130** | QH_4.5 | QH | 4.5 | | 0.991 | 0.972 | 0.000 | 0.011 | 0.000 | 0.250 | 0.083 | 0.625 | 0.229 | 168.604 | 16.350 | 0.000 | 3 | 37 |
| **131** | LPQ_4.5 | LPQ | 4.5 | | 0.978 | 0.974 | 0.000 | 0.005 | 0.000 | 0.125 | 0.063 | 0.375 | 0.229 | 176.614 | 24.360 | 0.000 | 2 | 81 |
| **132** | LPH_4.5 | LPH | 4.5 | | 0.978 | 0.973 | 0.000 | 0.007 | 0.000 | 0.250 | 0.083 | 0.500 | 0.167 | 183.634 | 31.380 | 0.000 | 3 | 102 |
| **133** | LQH_4.5 | LQH | 4.5 | | 0.991 | 0.971 | 0.000 | 0.011 | 0.000 | 0.250 | 0.083 | 0.625 | 0.229 | 168.604 | 16.350 | 0.000 | 3 | 38 |
| **134** | PQH_4.5 | PQH | 4.5 | | 0.984 | 0.974 | 0.000 | 0.008 | 0.000 | 0.250 | 0.083 | 0.500 | 0.167 | 177.741 | 25.487 | 0.000 | 3 | 89 |

| | | | | | | | | | | | | | | | | |
|---|---|---|---|---|---|---|---|---|---|---|---|---|---|---|---|---|
| **135** | LPQH_4.5 | LPQH | 4.5 | 0.984 | 0.973 | 0.000 | 0.009 | 0.000 | 0.250 | 0.083 | 0.500 | 0.167 | 177.741 | 25.487 | 0.000 | 3 | 90 |
